# A myelinic channel system for motor-driven organelle transport

**DOI:** 10.1101/2024.06.02.591488

**Authors:** Katie J. Chapple, Tabitha R.F. Green, Sarah Wirth, Yi-Hsin Chen, Ulrike Gerwig, Marie Louise Aicher, Yeonsu Kim, Lina Komarek, Angus Brown, Colin L. Crawford, Rebecca Sherrard Smith, Jeff Lee, Luis Pardo-Fernandez, Rebecca E McHugh, Celia M. Kassmann, Hauke B. Werner, Ilan Davis, Matthias Kneussel, Euan R Brown, Sandra Goebbels, Klaus-Armin Nave, Julia M. Edgar

**Affiliations:** School of Infection & Immunity, College of Medical Veterinary and Life Sciences, University of Glasgow, Glasgow, G12 8TA; Dept of Neurogenetics, Max Planck Institute for Multidisciplinary Sciences City Campus, Hermann-Rein-Strasse 3, D-37075 Goettingen, Germany; School of Life Sciences, University of Nottingham, University Park, Nottingham NG7 2UH; 4. School of Molecular Biosciences, College of Medical Veterinary and Life Sciences, University of Glasgow, Glasgow, G12 8TA; Department of Molecular Neurogenetics, Center for Molecular Neurobiology, ZMNH, University Medical Center Hamburg-Eppendorf, 20251, Hamburg, Germany; School of Engineering and Physical Sciences, Institute of Biological Chemistry, Biophysics and Bioengineering, Heriot Watt University, Edinburgh EH14 4AS, UK

**Keywords:** microtubule-dependent transport, live imaging, ageing, neurodegeneration

## Abstract

Myelin sheaths comprise compacted layers of oligodendroglial membrane wrapped spirally around axons. Each sheath, if imagined unwrapped, has a cytoplasm-filled space at its perimeter, linking it to the oligodendrocyte soma via a short process. By electron microscopy (EM), this space, which we term the ‘*myelinic channel system*’ contains microtubules and membranous organelles, but whether these are remnants of development or serve a function is unknown. Performing live imaging of myelinating oligodendrocytes expressing fluorescent reporters, we found that the myelinic channel system serves microtubule-dependent organelle transport. Further, the intra-myelinic movement of peroxisomes was modulated by neuronal electrical activity in these mixed neural cell cultures. Loss of oligodendroglial Kif21b or CNP *in vivo* led to apparent stasis of myelin organelles and secondary axon pathology. This suggests that oligodendrocytes require motor transport in myelin to maintain axonal integrity.

## Introduction

In the nervous system, the unusual size and architecture of neurons with long axons requires an intracellular transport system for moving cargo between the somata and distant axonal terminals. Axonal transport has been studied in detail since the 1960s (Allen et al., 1982; Austin et al., 1966; Brady et al., 1982; Grafstein, 1967; Lasek, 1967; Livett et al., 1968; Lubinska, 1964; Ochs et al., 1967; Weiss and Holland, 1967), including its molecular mechanism involving motor protein transport on microtubular tracts (Brady, 1985; Cason and Holzbaur, 2022; Hirokawa et al., 2010; Vale et al., 1985; Vargas et al., 2022; Zahavi and Hoogenraad, 2021). Here we show that a related logistical challenge exists at a much smaller scale with respect to the vital interaction of process-bearing oligodendrocytes and the axons they myelinate. Compacted myelin decreases the capacitance and increases the electrical resistance across the myelinated fibre (Cohen et al., 2020; Hartline and Colman, 2007). However, in so doing, myelin separates spatially the oligodendrocyte soma from the internodal axon-glial junction (Fig. 1A), a 20 nm wide synapse-like gap (Stys, 2011) thought to mediate the bi-cellular exchange of metabolites through monocarboxylate transporters (Fünfschilling et al., 2012; Lee et al., 2012).

**Fig. 1.**
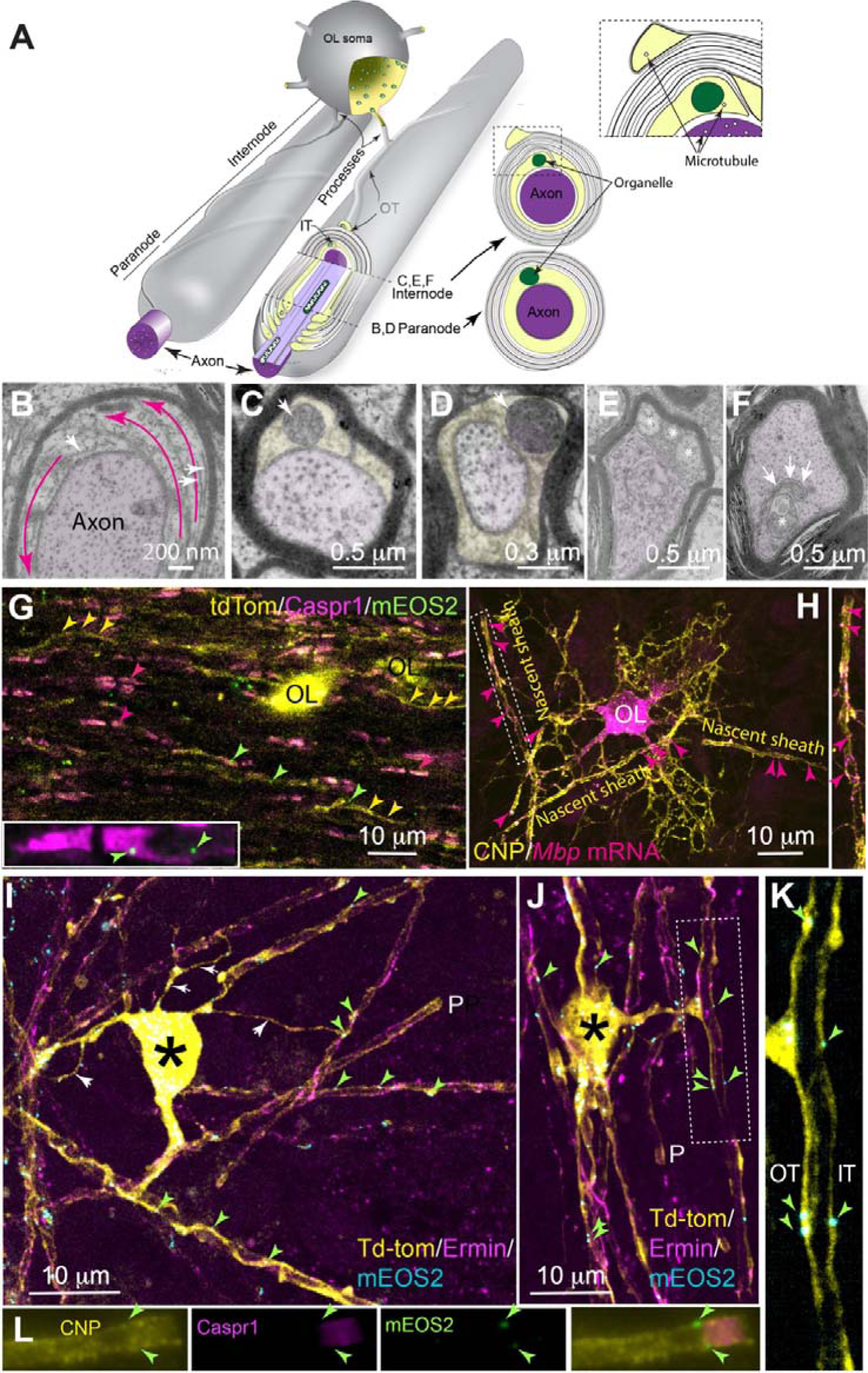
Microtubules and membranous organelles in the myelinic channel system. **A.** Schematic depiction of an oligodendrocyte (OL) soma and two myelin sheaths. Cytoplasm (yellow) fills the myelinic channel system including inner and outer tongues (IT, OT) that entwine the myelin sheath on its inner and outer surfaces, respectively, and the paranodal loops that are tethered to the axon at the paranode. Myelin organelles are represented in green. Two dashed lines depict approximately where the cross sections in the corresponding EM images are located. **B-F.** Micrographs of myelinated axon cross-sections including paranodal loops (B, D), the inner tongue (C, E), and an invagination of myelin membrane into the axon (F). In B, the white arrows indicate microtubules in paranodal loops, which encircle the axon as indicated by magenta arrows. In C, a multivesicular body occupies the inner tongue and in D, a dense organelle resembling a peroxisome resides in a paranodal loop. In E, three membrane-bound structures reside in myelin’s inner tongue. These have a pale core (asterisks), similar to the axoplasm, suggesting they are axonal sprouts. In F, an invagination of the glial cell membrane (arrows) surrounds an abnormal-appearing axonal organelle (asterisk), as shown previously in the PNS (Spencer and Thomas, 1974). **G.** Maximum intensity projection (MIP) from a confocal z-stack of a longitudinal section of the optic nerve in which oligodendrocytes and myelin are labelled with tdTomato (pseudo coloured yellow), oligodendrocyte peroxisomes are labelled with mEOS2 and paranodes are immunostained with anti-Caspr1, which is located on the axon. Peroxisomes are observed in somata (in this case, obscured by tdTomato) and myelin sheaths (yellow arrows), including paranodes (magenta arrows). A higher magnification view of individual channels in shown in **Supplementary Fig. 1M**. are shown Some peroxisomes reside in non-tdTomato rendered cells. Inset: two Caspr1 stained paranodes separated by a node of Ranvier. By inference, the two peroxisomes are located in paranodal and internodal myelin, respectively. **H.** MIP from a confocal z-stack of an early myelinating oligodendrocyte *in vitro* in which *Mbp* mRNA is labelled using single molecule fluorescence in-situ hybridisation (smFISH) and CNP is labelled using immunocytochemistry. *Mbp* mRNA can be observed in the cell body (OL), processes and nascent myelin sheaths. The cytoplasm filled spaces that stain with anti-CNP are remodelled over time, giving rise to the myelinic channels system depicted in the other images in this figure. **I.** MIP of a myelinating oligodendrocyte *in vitro* labelled with tdTomato and anti-Ermin, a cytoskeletal molecule that first appears during late myelination stages (Brockschnieder et al., 2006). The soma is marked with an asterisk and a paranode with P. mEOS2-labelled peroxisomes can be observed in the processes (white arrowheads) and myelin sheaths (green arrows). **J.** MIP of a second tdTomato-rendered oligodendrocyte *in vitro* showing peroxisomes (green arrows) in myelin sheaths including in the triangular shaped thickening at the junction of the process and the sheath delineated by the dotted line. The individual channels are shown in **Supplementary Fig. 1N**. **K.** High magnification view of sheath delineated by dotted line in I. Peroxisomes reside in both outer and inner tongues (OT and IT). **L.** Paranodal region from an oligodendrocyte *in vitro* labelled with anti-CNP (myelin) and anti-Caspr1. Two mEOS2-labelled peroxisomes are visible; by inference, the lower one is located just inside the paranodal region, marked by axonal Caspr1 expression, and the other at the juxtaparanode, between the internode and the paranode.

In the central nervous system (CNS), each myelin sheath derives from a lamellipod that extends from the oligodendrocyte soma. Growth of the prospective sheath involves actin remodelling (reviewed in Brown and Macklin, 2020) at the lamellipod’s leading edge (future ‘inner tongue’), which advances in a myelin basic protein-(MBP-) dependent fashion around the axon, underneath accumulating numbers of membrane wraps (Nawaz et al., 2015; Snaidero et al., 2014; Zuchero et al., 2015). Simultaneously, the lamellipod grows along the axon’s length (Snaidero et al., 2014) in response to calcium signalling in the oligodendrocyte (Baraban et al., 2018; Krasnow et al., 2018; Iyer et al., 2024), requiring nucleation of microtubules (Fu et al., 2019). Membrane compaction occurs by fusion of opposing intracellular surfaces (Aggarwal et al., 2011; Harauz et al., 2009) combined with tethering of opposing extracellular surfaces of the wrapping lamellipod (García-Mateo et al., 2018), culminating in tightly wrapped concentric layers of membrane. A cytoplasm-filled space remains at the perimeter of the sheath, if imagined unwrapped, suggesting a direct connection to the intracellular contents of the cell soma (Butt and Ransom, 1989) via the cell process (Peters, 1960; Sternberger et al., 1978) (Fig. 1A).

This cytoplasm filled space forms the ‘inner and outer tongues’ at the innermost and outermost edges of the wrapped sheath, and arrays of ‘paranodal loops’ at either end (Fig. 1A). The latter are tightly tethered to the axon by septate-like junctions (Bhat, 2003). Other cytoplasm-filled spaces in CNS myelin include transient openings of the compacted myelin (Snaidero et al., 2017; Velumian et al., 2011), rare Schmidt-Lanterman incisure-like openings on large fibres (Blakemore, 1969) and other conformations not yet fully defined (Edgar et al., 2021; Weruaga-Prieto et al., 1996). Myelin’s cytoplasm-filled spaces are generated and maintained, at least in part, by 2^/^,3^/^-cyclic nucleotide 3^/^-phosphodiesterase (CNP), which limits the membrane ‘compacting’ function of MBP (Snaidero et al., 2017; Trapp et al., 1988).

This myelinic channel system contain microtubules, endoplasmic reticulum, vesicle-like structures, multivesicular bodies, peroxisomes and Golgi outposts (Edgar and Griffiths, 2013; Frühbeis et al., 2020; Nakamura et al., 2021; Richert et al., 2014; Stassart et al., 2018). Since all these compartments have been implicated in developmental myelination, it remains unknown whether they exist as remnants of myelinogenesis, captured in the mature sheath, or play a functional role. We previously speculated that in the adult, the myelinic channel system provides a route for soluble oligodendroglial metabolites to reach the glial-axonal junction (Edgar et al., 2009; Meschkat et al., 2020; Nave, 2010a; Saab et al., 2013). Furthermore, since active transport processes can be visualised in cytosolic spaces of flattened myelin-like sheets in oligodendrocyte monolayer cultures (Ainger et al., 1993; Carson et al., 1997; Kachar et al., 1986; Herbert et al., 2017, Song et al., 2003), we additionally hypothesized that myelinic channels might serve as a route for the movement of membranous cargo to the glial-axonal junction, analogous to axonal transport to synaptic terminals.

Here we demonstrate microtubule-dependent organelle transport in established myelin sheaths and show that ablation from oligodendrocytes of kinesin 21B (Kif21b), a processive motor and modulator of microtubule dynamics (Ghiretti et al., 2016; Muhia et al., 2016; van Riel et al., 2017), leads to late onset secondary axon degeneration *in vivo*. This coincides with reduced levels of monocarboxylate transporter 1 (MCT1) in biochemically enriched myelin. Consistent with this, CNP-deficient mice, that lack myelinic channel integrity (Snaidero et al., 2017) and exhibit axonal degeneration (Edgar et al., 2009; Lappe-Siefke et al., 2003), and similarly in aged mice with structural myelin defects, the level of MCT1 in myelin is markedly diminished. As MCT1 resides at the glial-axonal junction and is thought to mediate the transfer of monocarboxylates between oligodendrocytes and axons (Fünfschilling et al., 2012; Lee et al., 2012), our data suggest myelinic transport is vital for oligodendrocyte-mediated axon support.

## Results

### The myelinic channel system and its organelle content

To define a putative transport infrastructure in the myelinic channel system (yellow in Fig. 1A), we used transmission electron microscopy (EM) of adult mouse optic nerve cross sections. As reviewed previously (Edgar and Griffiths, 2013), some paranodal loops of myelin contained microtubules (Fig. 1B), which were also observed in inner and outer tongues (**Supplementary Fig. 1A** and B). Multivesicular bodies (Fig. 1C and see Frühbeis et al., 2020) and single membrane structures resembling peroxisomes (Fig. 1D) were found in inner and outer tongues, paranodal loops, and very rarely, in cytosolic pockets of otherwise compact myelin (**Supplementary Fig. 1C**). Some membrane-bound structures in inner tongues (Fig. 1E) and paranodal loops had a pale core similar to the axoplasm, suggesting these are invading axonal sprouts (Steyer et al., 2023). Of the 2348 optic nerve axon cross-sections examined, 0.82 % (± 0.45 SD; n = 4 optic nerves of 12-month-old mice) of inner tongue processes contained structures resembling peroxisomes or multi-vesicular bodies. Based on (i) EM tissue sections being ∼50 nm thick, (ii) the average optic nerve myelin sheath length being ∼76 μm (Young et al., 2013) and the assumption that (iii) organelles are distributed randomly and (iv) spherical in shape, being ∼0.3 μm in diameter (Fig. 1C and D), our data suggest that the inner tongue of each optic nerve myelin sheath contains ∼2 such organelles (as paranodal loops are encountered only infrequently in EM images, we were unable to independently determine their organelle content using this method). In addition to myelin organelles, we observed invaginations of glial membrane into the axon (Fig. 1F and **Supplementary Fig. 1D-F)**, similar to previous reports in peripheral nerves (Spencer and Thomas, 1974). Very rarely, the myelin membrane even formed fusion-like profiles with the peri-axonal space (**Supplementary Fig. 1G**), suggesting direct exchange of materials between the two compartments.

As glutamate, which is released from electrically active axons (Micu et al., 2016), enhances the release of exosomes from oligodendrocytes *in vitro* (Frühbeis et al., 2020), we next asked if axonal electrical activity influences the localisation of organelles to myelin’s inner tongue, which faces the axon (Fig. 1A). To address this, we used an *ex vivo* optic nerve preparation in which we could precisely control action potential firing using stimulating and recording suction electrodes (**Supplementary Fig. 1H**), as described (Stys et al., 1991). One of each pair of adult mouse optic nerves was unstimulated while the contralateral nerve was stimulated to fire constitutively at 7 or 50 Hz for 20 minutes. By EM of nerve cross sections, the stimulated nerve had more inner tongues with organelles (average 0.65% ± 0.28 SD: ∼1.7 organelles per sheath) compared to the unstimulated contralateral nerve (average 0.32 % of fibres ± 0.34 SD; ∼0.8 organelles per sheath), although with 5 nerve pairs analysed, the differences were not significant (**Supplementary Fig. 1I-K**). While organelle numbers could also decrease due to fusion, these data led us to hypothesize that organelle transport to or from the inner tongue might be modulated by axonal electrical activity.

Next, we focussed on peroxisomes using a transgenic mouse in which oligodendroglial peroxisomes are labelled with the photoconvertible fluorescent protein mEOS2 (Richert et al., 2014). To demarcate the myelinic channel system, we additionally labelled a proportion of oligodendrocytes with a transgenically-expressed, tamoxifen-inducible fluorescent protein tdTomato. In longitudinal sections of optic nerve, mEOS2-labelled peroxisomes were observed in the oligodendrocyte soma; at paranodes, as indicated by axonal expression of Caspr1; and in internodal myelin (Fig. 1G; **Supplementary Fig. 1M**). This was confirmed in spinal cord (**Supplementary Fig. 1L**). However, in the densely packed white matter it was difficult to resolve individual myelin sheaths, even with sparse labelling. Therefore, for subsequent observations, we used a murine spinal cord-derived myelin-forming co-culture of neurons and glia (“myelinating cell culture”), in which axons are enwrapped in compact myelin (Bijland et al., 2019; Thomson et al., 2008). In nascent sheaths, *Mbp* mRNA, which is translated locally in myelin (Wake et al., 2011) could be observed in the CNP +ve cytoplasm filled spaces that are eventually remodelled to form the myelinic channel system (Fig. 1H; **Supplementary Video 1**). In established sheaths, identified by intensely tdTomato positive paranodal regions, oligodendrocyte peroxisomes were observed in tdTomato rendered somata, processes, and sheaths (Fig. 1I), including inner and outer tongues (Fig. 1J and K; **Supplementary Fig. 1N**) and 48.2 % (± 7.4 SD) of paranodes examined (Fig. 1L; **Supplementary Fig. 1O**).

### Myelinic organelle transport is microtubule dependent

The optic nerve stimulation experiment suggests that myelin organelles are motile. To explore this further, we used live imaging of mEOS2+ oligodendrocyte peroxisomes in myelinating cell cultures in which oligodendrocytes wrap axons with appropriately thick compact myelin sheaths (Edgar et al., 2021; Thomson et al., 2008). As in other mammalian cell types (Wali et al., 2016) most myelin peroxisomes were apparently stationery or underwent only short-distance, Brownian-like motion. However, within a 10-minute imaging window, 16.2% (± 4.07 SD) of myelin peroxisomes exhibited movement (traversing distances ≧ 3 μm; Fig. 2A and B) and covering distances up to ∼70 µm. Travel was punctuated by periods of stasis (Fig. 2A), such that the overall velocity over 10 minutes was approximately 10× less than when motile; the latter being 0.18 µm/sec (± 0.02 SD), on average. Some peroxisomes travelled predominantly in one direction (Fig. 2A-C), while others reversed direction (indicated by cyan circles in **Supplementary Fig. 2A**) and/or crossed paths with another (indicated by orange crosses in **Supplementary Fig. 2A**). Often, peroxisomes in the outer tongue (where it could be determined definitively) travelled towards the paranode (Fig. 2B and C; **Supplementary Videos 2 and 3**). In **Supplementary Videos 4 and 5** a peroxisome travels to the paranodal loops of myelin via the presumptive outer tongue, traverses 4 paranodal loops and travels back in the direction of the cell body via the presumptive inner tongue.

**Fig. 2.**
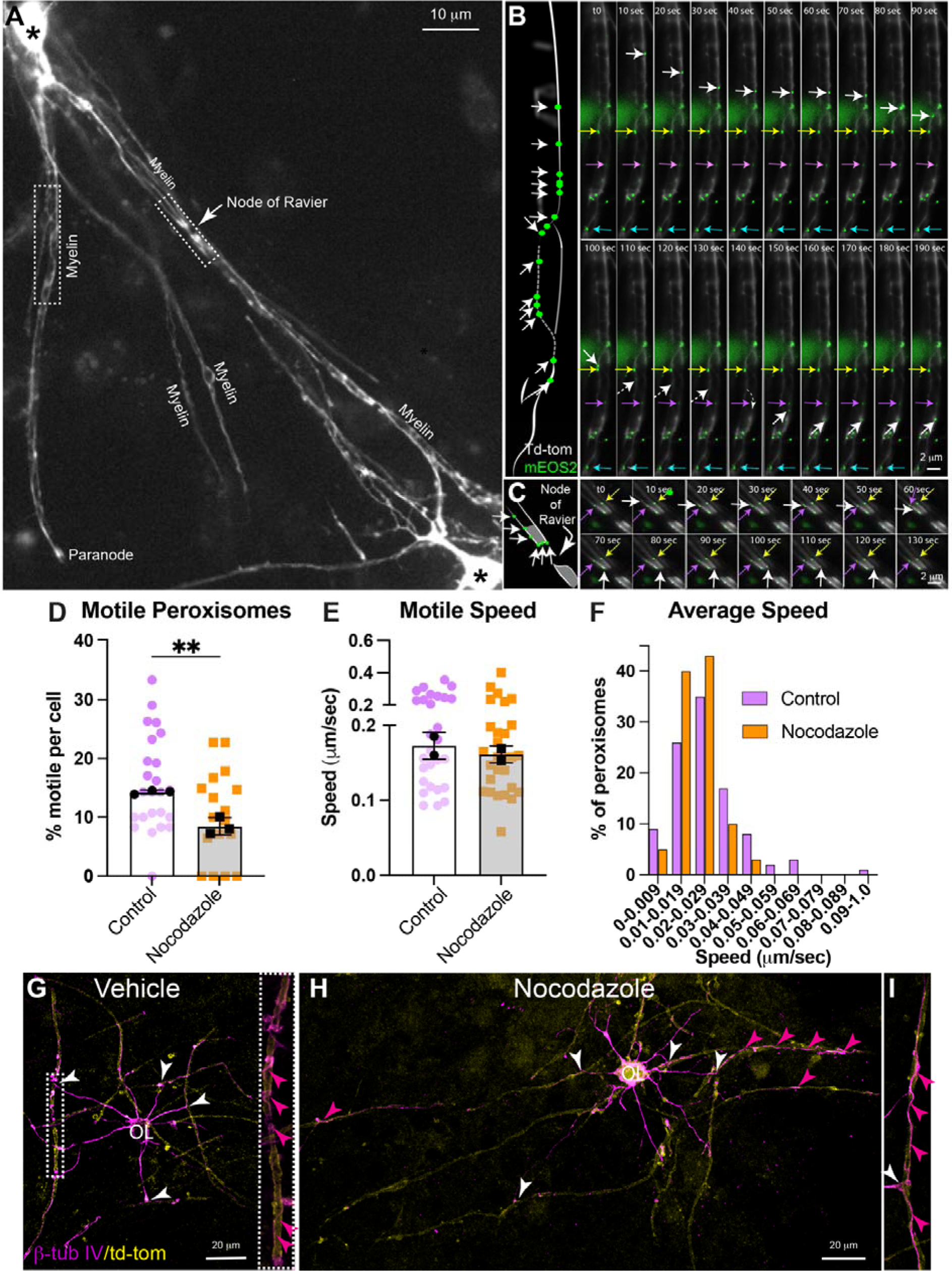
Myelinic peroxisomes transport is microtubule-dependent. **A.** Overview of two tdTomato rendered oligodendrocytes (somata are labelled with asterisks) from which the timelapse images in B and C were acquired. **B and C.** The schematics on the left illustrate the totality of the movement of a single myelin peroxisome in each of the timelapse movies depicted on the right. White lines represent cytoplasm-filled spaces within (solid lines) and outside (dashed lines) the focal plane. In B, the timelapse images show a motile peroxisome (white arrows) move down the presumptive outer tongue towards the paranodal loops (see overview), slowing or stalling at 30, 110 and 160 second time points. At 140 seconds, the organelle moves out of the focal plane and the path (small white arrow) is inferred from the movie. Most peroxisomes (coloured arrows) were stationery. In C, paranodal regions of adjacent sheaths, are separated by a node of Ranvier (see A). A motile peroxisome (white arrow) moves down the outer tongue process (determined from the relationship to the node of Ranvier) and traverses the presumptive paranodal loop nearest the node of Ranvier, before moving out of the focal plane. Other peroxisomes (coloured arrows) were stationary. **D.** Nocodazole (20 μM, 3-hour incubation) reduced the percentage of myelin peroxisomes that were motile during the 10-minute period. Coloured symbols represent individual oligodendrocytes (n = 22 and 17, control and nocodazole, respectively), black symbols represent experimental medians (n = 3 each control and nocodazole); bars indicate mean of the median values ± SD. An unpaired Student’s *t* test was used to compare the experimental means. ** *p* < 0.01. **E.** Nocodazole appeared not to alter the velocity of individual myelin peroxisomes when motile. Coloured symbols each represent one motile peroxisome (n = 28 and 29, control and nocodazole respectively) and black symbols represent experimental medians (n = 2 each, control and nocodazole); bars indicate the mean of the experimental median values ± SD. **F**. Nocodazole caused a shift to slower average velocities, reflecting motile and stationary phases, of individual peroxisomes during the 10-minute imaging period, likely reflecting a reduction in available transport tracks. **G-I.** β−tubulin IV staining of tdTomato labelled oligodendrocytes, following extraction of free tubulins (see also **Supplementary Video 6**), treated with DMSO (vehicle; G) or nocodazole (20 μM, 3 hours; H and I). White arrows indicate where processes from the oligodendrocyte attach to the myelin sheath and magenta arrows indicate regions of continuous β−tubulin IV staining. The sheath delineated by a white rectangle in G is shown at higher magnification in the inset (individual channels are shown in **Supplementary Fig. 2B**). The sheath in I is from a different cell from that in H (individual channels are shown in **Supplementary Fig. 2B**).

In other eukaryotic cell types, peroxisomes are transported on microtubules (Rapp et al., 1996 and reviewed in Covill-Cooke et al., 2021). To determine if this is also true in the myelinic channel system, we treated myelinating cell cultures with nocodazole, which interferes with the polymerization of microtubules (Kesarwani et al., 2020). As axons also contain microtubules, we aimed to minimise confounding effects on myelinic transport (caused by axon dysfunction) by using a concentration and incubation time that did not markedly alter the microtubule network in neurons. In these cell dense cultures, 3-hours in 20 μM nocodazole led only to minor alterations in anti-β-tubulin 3 staining (**Supplementary Fig. 2B**), indicating that neuronal microtubules were only partially disrupted over this time period. Furthermore, detyrosinated tubulin, a non-cell type specific marker of stable microtubules, appeared similar to controls (**Supplementary Fig. 2B**). Using this protocol, nocodazole caused a significant reduction in the percentage of motile myelinic peroxisomes (Fig. 2D) without altering their velocity during motile phases (Fig. 2E). Nonetheless, the average velocity of individual organelles (including motile and stationary phases) within the 10-minute imaging period, shifted towards slower speeds (Fig. 2F). Consequently, the maximum distance travelled by a single peroxisome in 10 minutes in this experimental series was reduced from 55.0 μm in controls to 23.8 μm in nocodazole-treated cells, likely reflecting the drug’s perturbation of microtubular tracks. Indeed, in DMSO-treated control cultures, β-tubulin IV (which is specific in the CNS to oligodendroglia) stained continuous structures within myelinic channels in 94 % (± 2.0 SD) of tdTomato positive cells (Fig. 2G, **Supplementary Fig. 2B, Supplementary Video 6**; *n* = 33 cells; 2 independent cell cultures), but in only 45 % (± 7.0 SD) of tdTomato positive cells after nocodazole-treatment (Fig. 2H, **Supplementary Fig. 2B**; *n* = 17 cells; 2 independent cell cultures). Notably, in both control and nocodazole-treated cultures, the amount of polymerised β-tubulin IV varied considerably from sheath to sheath and from cell to cell. In summary, myelinic channels enable an active, microtubule-dependent transport of peroxisomes, and presumably other membrane-bound cargo.

### Myelin microtubules harbour post-translational modifications

Our data suggested that some myelin microtubules are more resistant to nocodazole than others, suggesting that myelin contains mixed populations of “dynamic” and “stable” microtubules (Jansen et al., 2023). Microtubule stability is encoded in post-translational modifications (PTMs) of tubulins, and the presence of differentially modified tubulins is crucial in the assembly, disassembly and rearrangement of the microtubule cytoskeleton (Borys et al., 2020; Janke and Bulinski, 2011). Recently, genetic sensors were developed that identify tyrosinated microtubules (AlaY1; Kesarwani et al., 2020), which are generally considered to be rather labile; or stable microtubules (StableMARK; Jansen et al., 2023), which are generally acetylated and/or detyrosinated. We observed both genetic sensors in oligodendrocyte cell bodies, processes, and myelin sheaths (Fig. 3A and B). However, as StableMARK can by itself stabilise microtubules when overexpressed, and as AlaY1 also labels free tubulins, we next used antibody staining, following extraction of free tubulins (Jansen et al., 2023), to identify PTMs on microtubule polymers. Anti-tyrosinated tubulin partially labelled one or more myelin sheaths in 83% of all tdTomato positive oligodendrocytes (*n* = 12 cells; 3 independent cell cultures) (Fig. 3C and D; **Supplementary Fig. 3**). Anti-detyrosinated tubulin was not observed in myelin sheaths (Fig. 3E; **Supplementary Fig. 3**). Acetylated tubulin was observed in some sheaths (Fig. 3F and G; **Supplementary Fig. 3**), as reported *in vivo* previously (Kusch et al., 2017; Werner et al., 2007). Because the antibodies to tyrosinated and acetylated tubulin are raised in the same species, we were unable to determine if different PTMs co-exist in the same sheaths.

**Fig. 3.**
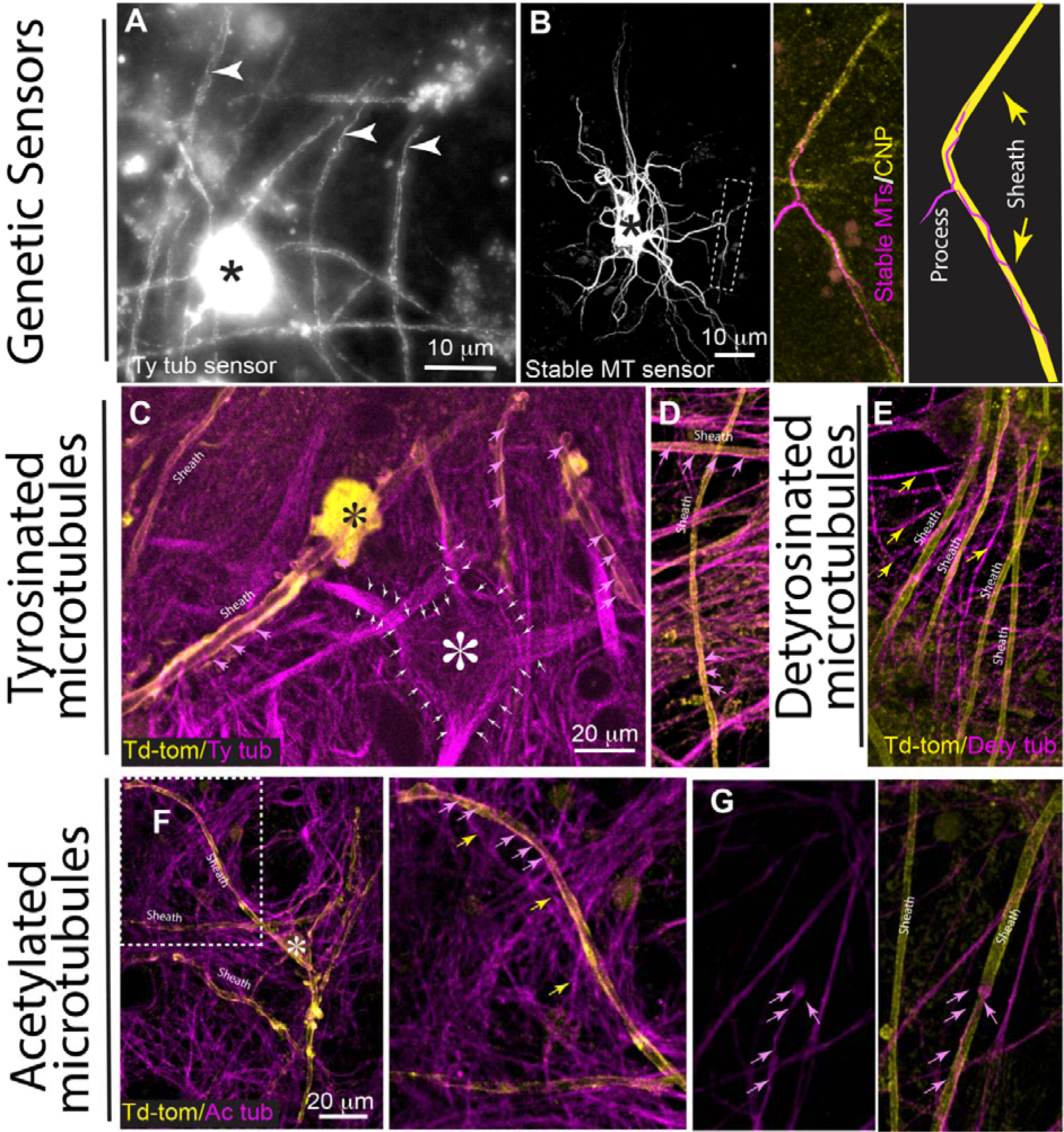
Microtubules in the myelinic channel system harbour various post-translational modifications. **A** and **B**. Nine hours after oligodendrocyte transfection, genetic sensors labelling tyrosinated tubulins (Ty tub; A) and stable microtubules (stable MTs; B) in myelin, were visible. In A, arrowheads point to continuous-appearing structures suggesting they are microtubule polymers. In B, the cell body labelling is saturated, and processes are intensely labelled therefore a higher magnification view of an associated myelin sheath (delineated by the broken line), is shown on the right, stained with anti-CNP and illustrated for clarity in the schematic. The abnormal appearance of the somata and processes is likely because StableMARK itself stabilises microtubules if overexpressed. **C-F.** Following extraction of free tubulins, immunocytochemistry was used to identify PTMs on myelin microtubules. Anti-tyrosinated tubulin labelled some myelin sheaths (magenta arrows) of tdTomato labelled cells, and the cell bodies (asterisk) and processes of other cell types (C). In general, anti-tyrosinated tubulin only partially labelled sheaths (magenta arrows; D). Anti-detyrosinated tubulin did not colocalise with tdTomato positive sheaths but labelled other cell processes (yellow arrows). Where magenta and yellow coincide, the tubulin stain does not follow the contours of the sheath, suggesting it is the processes of overlapping cells (E). Anti-acetylated tubulin labelled a few myelin sheaths (magenta arrows) although it was prevalent in other cellular processes (yellow arrows). The oligodendrocyte soma is labelled with an asterisk in the overview image on the left (F). Anti-acetylated tubulin only partially labelled sheaths (G). Individual channels from C, D, E and F are shown in **Supplementary Fig. 3**, for clarity.

Together these data demonstrate tubular tracks within the myelinic channel system are required for active translocation of organelles. Interrogation of our databases of purified myelin proteins, identified by label-free mass spectrometry (Gargareta et al., 2022; Jahn et al., 2020), provided evidence for the presence in myelin of molecular motor proteins, including dynein and kinesin (**Supplementary Table 1A**). Tubulins, microtubule binding proteins and tubulin modifiers were also identified in biochemically isolated ‘myelin’ (**Supplementary Table 1B**).

### Modulators of neuronal activity influence the motility of myelin peroxisomes

Myelin-associated peroxisomes are essential for the long-term integrity of axons, particularly faster spiking axons, which are most vulnerable to peroxisomal defects (Kassmann et al., 2007). Furthermore, axo-myelin glutamatergic signalling and axonal electrical activity trigger neuroprotective exosome release and oligodendroglial metabolic support (Frühbeis et al., 2020; Saab et al., 2016). As a simple cellular model to examine the response of myelin-associated peroxisomes to changes in neuronal firing, we used myelinating cell cultures in which neuronal electrical activity and neuronal energy consumption can be enhanced with the GABA receptor blocker picrotoxin (PTX; 100 μM) or diminished with tetrodotoxin (TTX; 1 μM) (Fig. 4A and B).

**Fig. 4.**
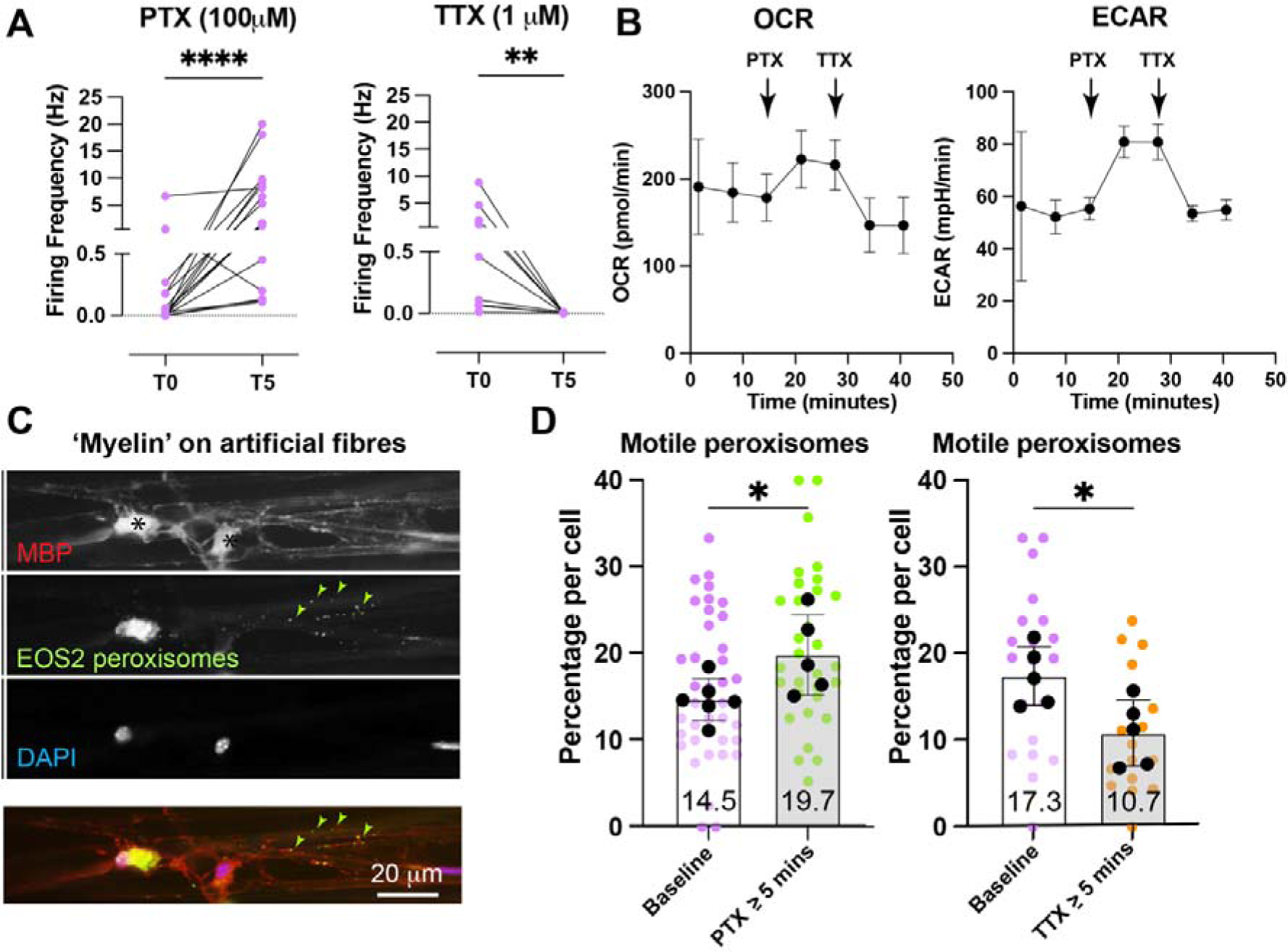
Neuronal activity locally modulates the movement of myelin peroxisomes. **A.** In myelinating cell cultures, PTX or TTX enhance or block neuronal activity, respectively. Graphs show firing frequency of individual neurons recorded in current clamp at baseline (T0) and 5 minutes (T5) after the application of drug. A paired Student’s t test was used to compare pre- and post-drug values. Results are from 9 and 5 independent cell cultures, respectively **** p = <0.0001; ** p = <0.01. **B.** In neuron-enriched cultures, oxygen consumption rate (OCR) and extracellular acidification rate (ECR) changed in response to PTX or TTX indicating a general increase and decrease, respectively, in energy consumption. Symbols represent the mean of 3 independent experiments ± SD. **C.** Widefield epifluorescence micrograph of peroxisomes in the MBP positive myelin-like structures formed on inert fibres, demonstrating that neuronal activity is not necessary for peroxisomes to enter the sheath during myelinogenesis. **D.** Graphs showing the proportion of myelin peroxisomes per cell that are motile during a 10-minute imaging period. Coloured dots represent individual cells and black dots represent experimental medians. Bars represent mean of experimental medians ± SD. PTX or TTX significantly increased or decreased, respectively, the percentage of myelin peroxisomes that were motile during a 10-minute imaging period. An unpaired Student’s t test was used to compare pre- and post-drug values between the median values of the independent experiments. Results are from 5 or 6 independent cell cultures. *p = <0.05.

Live imaging of oligodendrocyte processes over a 10-minute timeframe (beginning 5-minutes post drug application) showed that modulating neuronal activity had no effect on retrograde or anterograde transport of peroxisomes between the myelin sheath and the oligodendrocyte soma (**Supplementary Fig. 4A, Supplementary Video 7**). This concurs with our observation that peroxisomes entered the ‘myelin sheath’ even when formed on inert fibres (Fig. 4C). Nonetheless, in the presence of neurons, myelin-associated peroxisomes responded to PTX or TTX by becoming more or less motile, respectively (Fig. 4D), demonstrating that neuronal activity influences myelinic transport processes locally. Despite this, the number of peroxisomes at paranodes, where myelin is tethered to the axon, did not change in response to changes in neuronal activity, at least in the short term (**Supplementary Fig. 4B**).

### Deleting Kif21b from oligodendrocytes leads to secondary axon loss

In the CNS, β-tubulin IV is specific to oligodendrocytes and localises to the somata, processes and myelin (Edgar et al., 2021; Terada et al., 2005 and Fig. 3 this manuscript), leading us to hypothesize that β-tubulin IV could play a role in cargo transport, sheath formation and axon maintenance. However, targeting the *Tubb4* gene selectively in oligodendrocytes in *Tubb4^flox/flox^*::*Cnp1*^+/Cre^ mice did not lead to overt changes in myelinated CNS fibres, even at 14 months of age, except that the g-ratio tended to be slightly higher than in controls (**Supplementary** Fig.5). Thus, *Tubb4* function is likely compensated for by other β-tubulins.

Next, we turned to a motor protein, Kif21b that we identified as a putative candidate for a function in the myelinic channel system because polymorphisms in *KIF21B* have been associated with increased susceptibility to multiple sclerosis (MS). Moreover, *KIF21B* is upregulated ∼10 fold in MS white matter irrespective of genotype (Goris et al., 2010; Kreft et al., 2014) and expression of *Kif21b* is differentially regulated in different populations of mature oligodendrocytes in spinal cord of experimental autoimmune encephalomyelitis (Zheng et al., 2023). Kif21b is both a plus end directed progressive motor protein and a modulator of microtubule dynamics (Ghiretti et al., 2016; Hooikaas et al., 2020; Muhia et al., 2016; van Riel et al., 2017) that in neurons, is enriched in dendrites and growth cones at neurite tips (Labonté et al., 2014; Marszalek et al., 1999). To determine if Kif21b is expressed in oligodendrocytes in the optic nerve, we compared by western blotting, optic nerve lysates from *Kif21b* conditional knockout (c*Kif21b* KO) mice (*Cnp1^Cre^*^/+^::*Kif21b*^flox/flox^) with two controls (*Cnp1^+^*^/+^::*Kif21b*^flox/flox^ and *Cnp1^+^*^/Cre^::*Kif21b*^+/flox^ ^or +/+^). Whilst *Cnp1^Cre^* can recombine ectopically (Genoud et al., 2002), retinal ganglion cells (whose axons traverse the optic nerve) do not recombine, and in the optic nerve, all recombined cells are oligodendrocytes (Saab et al., 2016; Späte et al., 2024). Kif21b was not detected by western blotting of whole optic nerve lysates from c*Kif21b* KO mice but was readily detectable in control optic nerves (Fig. 5A), demonstrating in this white matter tract, Kif21b is predominantly expressed by *Cnp1*-expressing oligodendrocytes.

**Fig. 5.**
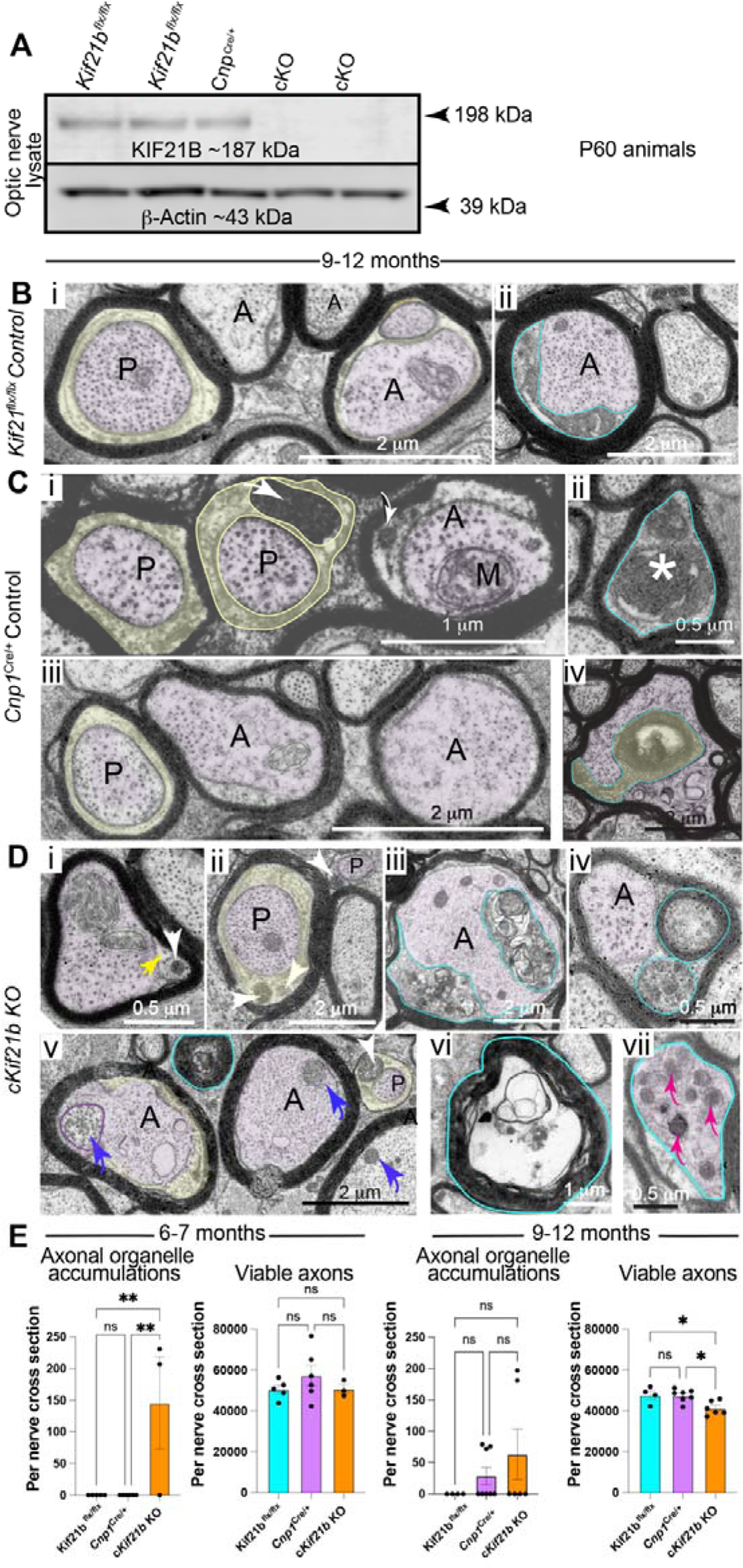
Depletion of Kif21b from oligodendrocytes leads to secondary axon loss. **A.** In the optic nerve, a pure white matter tract lacking neuronal cell bodies and dendrites, *Cnp1*Cre-dependent deletion of *Kif21b* led to depletion of Kif21b in whole nerve lysate, demonstrating this motor protein is expressed in oligodendrocytes **B.** Electron micrographs of optic nerve axons from 9-12 month *Cnp1*^+/+^::*Kif21b*^flx/flx^ controls. Note the exceptional finding of periaxonal electron dense material (blue outline, Bii). **C.** In *Cnp1*^Cre/+^ control optic nerves, myelinated axons appeared largely normal. Periaxonal electron dense organelles in Ci (white arrows) are also seen in WT mice (Fig. 1). Rarely, axons appeared electron dense and degenerating (asterisk Cii) or myelinic channels were abnormally extended (blue outline in Civ). **D.** In *Kif21b* conditional KO nerve (c*Kif21b* KO), most myelinated axons appeared morphologically normal (Di and ii). Myelinic organelles were observed in myelin’s inner tongue (Di) and paranodal loops (Dii) as in WT (Fig. 1). In Di, the yellow arrow points to a microtubule. However, myelinic channels were occasionally locally enlarged, impinging on the axon (delineated in blue in Diii). Redundant myelin and/or abnormal electron dense material were observed in some swollen inner tongues (delineated in blue in Div). Extracellular myelin whorls (blue outlines in Dv and vi) were not more frequent than in controls. Rarely, axons contained abnormal-appearing mitochondria (blue arrow on the left in Dv in comparison to the two mitochondria indicated by blue arrows on the right). Axonal organelle accumulations were also a feature of mutant mice (magenta arrows in Dvii.) **E.** Quantitation of axonal organelle accumulations and total axon numbers in c*Kif21b* KO optic nerves and controls at age 6-7 (left) and 9-12 months (right). Data were analysed by one-way ANOVA followed by Tukey’s multiple comparisons test. * p < 0.05; ** p < 0.01. A: axon; P: paranode.

To determine if ablating Kif21b from oligodendrocytes had consequences for the myelinated axon, we examined optic nerve cross sections by EM (Fig. 5B-D). Across all genotypes, most myelinated axons appeared morphologically normal at 1 year of age (Fig. 5Bi; 5C i,iii and 5D i, ii). However, in c*Kif21b* KO mice, inner tongues often appeared distended and contained membranous material (Fig. 5D iii, iv) and occasionally otherwise healthy-appearing axons contained mitochondria lacking normal-appearing cristae (Fig. 5D iv). Accumulations of axonal organelles (magenta arrows, Fig. 5D vi), indicative of axonal transport stasis (Edgar et al., 2004), were observed in c*Kif21b* KO mice from 6 months of age (Fig. 5E). Importantly, these pathological features were seen also at 4 months of age but largely “lost” by age 9-12 months; the decline in absolute axon number by 9-12 months suggesting these pathologically affected axons had degenerated at this age (Fig. 5E).

While similar pathological axonal changes were seen occasionally also in heterozygous *Cnp1*^+/Cre^ controls at 9-12 months of age, there was no comparable loss of optic nerve axons suggesting that Kif21b-dependent transport in oligodendrocytes (and/or microtubule destabilisation) is required for long-term axonal integrity.

### A role for CNP1 and Kif21b in maintaining monocarboxylate transporter levels

We hypothesized that axonal integrity is impaired when metabolic support by oligodendrocytes is compromised. Transport processes in myelinic channels could contribute to overall myelin protein turnover, including that of MCT1 at the glial-axon junction. To determine if oligodendroglial Kif21b depletion impacts specific protein levels in myelin, we performed western blotting of spinal cord myelin from mice at postnatal day 60 (P60) and 1 year. As expected, Kif21b itself was strongly reduced in purified myelin from c*Kif21b* KO mice (Fig. 6A), and to a lesser degree in total spinal cord homogenate containing both white and grey matter (**Supplementary** Fig.6). However, the *Cnp1^Cre^* knock-in allele (resulting in reduced *Cnp1* gene dosage) was likely to contribute to these effects as CNP acts as a strut that maintains the integrity of myelinic channels (Snaidero et al., 2017) and *Cnp1* heterozygosity leads to myelin dysfunction at advanced age (Hagemeyer et al., 2012). Indeed, in myelin from 1 year old *Cnp1*^+/Cre^ littermate controls, Kif21b levels became remarkably variable (Fig. 6A). For comparison, myelin oligodendrocyte glycoprotein (MOG), a transmembrane protein on the outer myelin layer, was unaltered in c*Kif21b* KO mice compared to either control. In contrast, levels of MCT1, which is required at the glial-axonal junction for the transfer of pyruvate/lactate to the periaxonal space (Fünfschilling et al., 2012; Lee et al., 2012) were significantly reduced in 1 year old mutants, but not yet in their P60 counterparts (Fig. 6A). Indeed, MCT1 was also reduced in 1 year old *Cnp1^+/Cre^* controls, confirming that both Kif21b and CNP are required for maintaining normal MCT1 levels at the glial-axon junction. Importantly, there was no significant change in the levels of β-tubulin IV in purified myelin from c*Kif21b* KO mice, suggesting there was no lack of tubular tracks. We thus hypothesize that the loss of myelinic channel integrity (with reduced CNP1 and absence of oligodendroglial Kif21b) causes a slow attrition in delivery of MCT1 to the glial-axonal junction.

**Fig. 6.**
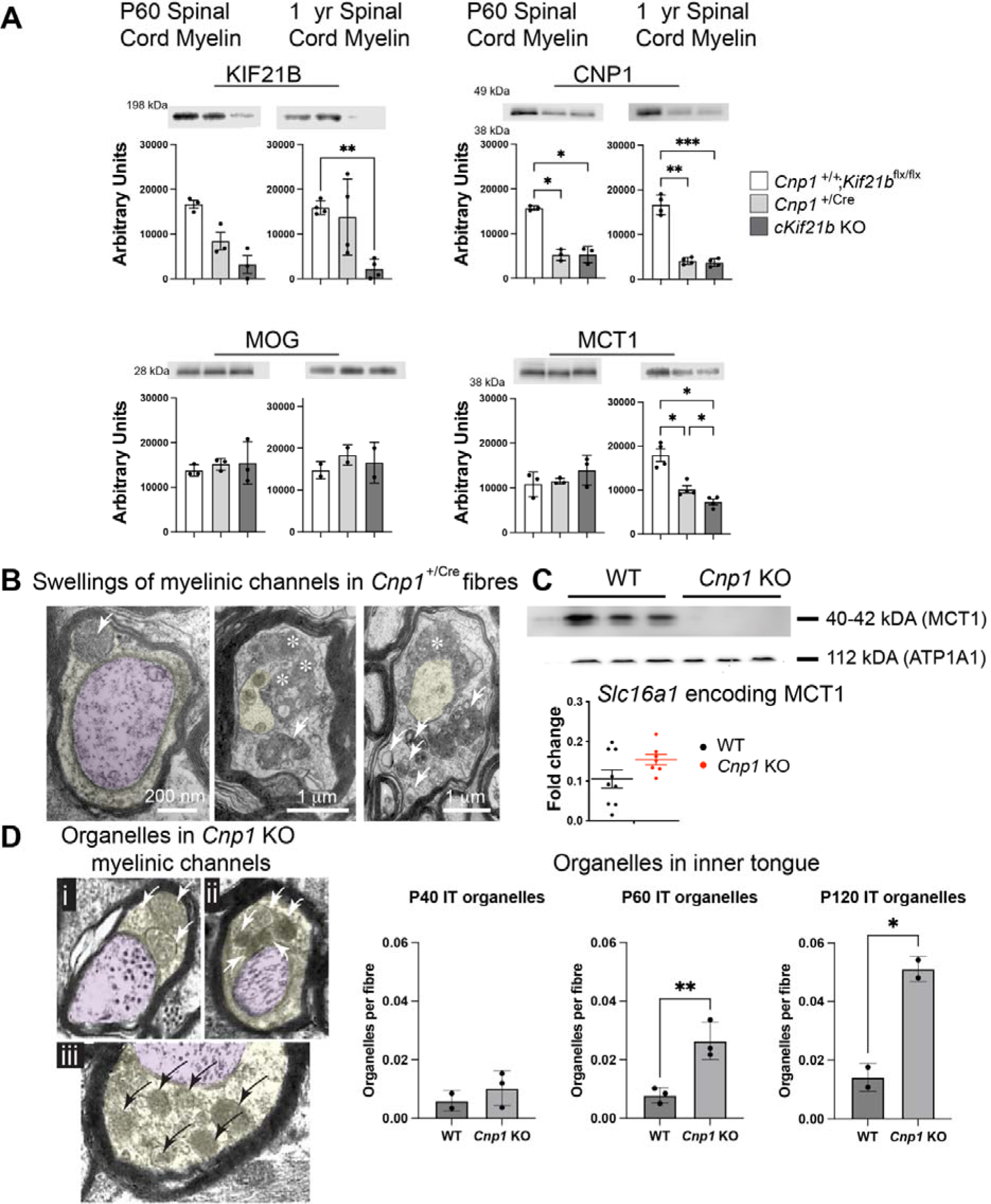
Monocarboxylate transporter 1 is reduced when myelinic transport is impaired. **A.** Western blots of spinal cord myelin-enriched fractions of P60 and 1 year old mice. Levels of Kif21b were reduced in *Kif21b* conditional knockout (c*Kif21b* KO) mice, albeit that the reduction was not significant at P60 due to intra-group variation. Unexpectedly, levels in the *Cnp1*^Cre/+^ controls (Cre cont) were highly variable. Levels of CNP1 were significantly reduced in both the *Cnp1*^Cre/+^ and c*Kif21b* KO mice compared to the *Cnp1*^+/+^::*Kif21b*^flx/flx^ (WT) control, at P60 and 1 year. Levels of MOG were similar in all three genotypes, at both ages. At P60, MCT1 levels were similar in all three genotypes, but at 1 year of age, levels were significantly reduced in the *Cnp1*^Cre/+^ mouse compared to WT control and in the c*Kif21b* **KO** compared to both WT and *Cnp1*^Cre/+^ controls. Data were analysed using a one-way ANOVA followed by Tukey’s multiple comparisons test. *: p < 0.05; **: p < 0.01; ***: p<0.001 **B.** Electron micrographs of myelinated optic nerve axons from heterozygous *Cnp1*^+/Cre^ mice at age 14 months. On the left, a normal appearing paranode containing a peroxisome-like organelle (white arrow). On the right, membranous structures (arrows) and dense cytoplasm (asterisks) appear to accumulate in inner tongues in a small number of fibres. Inner tongues appear grossly swollen and impinge upon the axon. **C.** Western blotting of myelin purified from homozygous *Cnp1* knockout mice shows a dramatic reduction in steady state levels of MCT1. In contrast, RT-PCR of brain lysates demonstrates no significant change in *Slc16a1* mRNA levels, when examined in relation to *Slc16a1* mRNA levels in myelin from P20 WT mouse brain, in agreement with a post-transcriptional MCT1 deficit. Data were analysed using a two tailed, unpaired Student’s t test. **D. Left**: Electron micrographs of myelinated optic nerve axons from homozygous *Cnp1* knockout mice, taken at (*i*) P40, (*ii*) P60 and (*iii*) P120, with undefined organelles (arrows) residing in the inner tongues (yellow overlay). **Right:** Quantitation demonstrates these organelles accumulate from age P60, i.e. long before clinical symptoms emerge at age 6 months (Lappe-Siefke et al., 2003). Bars represent mean ± SD. Data were analysed using a two tailed, unpaired Student’s t-test. Data points represent mean values from electron micrographs of optic nerves of individual animals. *: p < 0.05; **: p < 0.01.

### The role of CNP1 in organelle transport through myelinic channels

To test our hypothesis further, we compared myelin composition and myelinated fibre morphology in heterozygous (*Cnp1*^+/Cre^) and homozygous (*Cnp1*^-/-^) mutants. In optic nerves from *Cnp1* heterozygous mice, at older age (14 months), membranous structures (arrows) and dense cytoplasm (asterisks) were occasionally observed in enlarged inner tongues (Fig. 6B), resembling changes in the aged CNS (Peters and Kemper, 2012). In homozygous *Cnp1* mutants, with myelinic channel collapse (Snaidero et al., 2017), axonal changes in myelinated fibres begins as early as P5 (Edgar et al., 2009) and likely underlie premature death, which occurs before one year of age (Lappe-Siefke et al., 2003). When analysing homozygous *Cnp1* mutant mice by western blotting, we found that MCT1 was strongly reduced in a myelin-enriched brain fraction by at least P75 while transcription of the encoding gene (*Slc16a1*) was similar in mutant and control brains (Fig. 6C). This suggests the absence of CNP affects the delivery of MCT1 to the glial-axonal junction and/or its turnover. In support of impaired intra-myelinic transport, we found by EM of optic nerve, that in some fibres in homozygous *Cnp1* mutants, organelles appeared trapped within the myelinic channel system (Fig. 6D).

### White matter ageing is associated with impaired intra-myelin transport

Advanced age is associated with altered structural integrity of myelin in white matter tracts and intracortical myelinated fibres (Mueller, 2024; Peters, 2002), and with axon vulnerability as assessed by magnetic resonance imaging (MRI) in humans (Burzynska et al., 2024). We hypothesised that myelin changes include alterations in the myelinic channel system, similar to its destabilization by loss of CNP. This is suggested by a peculiar interaction of *Cnp1* heterozygosity and advanced brain age on behavioural symptoms in mice (Hagemeyer et al., 2012). To directly determine if age-related white matter changes impact intra-myelinic transport processes, we compared myelin morphology and biochemistry in wildtype mice at age 6 and 24 months. We first measured myelin inner tongue sizes across axon diameters, observing a volume increase in the aged mice, most notably in small diameter fibres. This was accompanied by a twofold increase in fibres with organelles in the inner tongue (Fig. 7A). Furthermore, these changes were observed in the presence of an almost two-fold reduction in the overall density of myelinated axons at 24 months (corpus callosum) and a doubling of axonal degeneration profiles (Fig. 7B), raising the possibility that morphologically more severely affected fibres had been lost at this age.

**Fig. 7.** White matter aging leads to myelinic channel changes, decreased MCT1 levels and axon degeneration. **A.** Electron micrograph of optic nerves from a 24-month-old mouse preserved by high pressure freezing (HPF), demonstrating that inner tongues (highlighted in yellow) are occasionally enlarged and are more likely to contain electron dense organelles (black arrowheads). The graphs on the right shows that inner tongue areas tended to be increased in 24-month-old animals compared to 6-month-old mice across all axon diameters (180-230 axons per nerve were examined), reaching significance in smaller diameter fibres. Data were analysed using multiple unpaired t-tests, corrected for multiple comparisons using Holm-Šídák method (n = 5 or 6; * p<0.05). A higher proportion of myelinated fibres have organelles in the inner tongue (IT; n = 3). Data were analysed using an unpaired two-tailed Student’s t test. * p < 0.05. **B.** HPF EM overview showing that in 24-month-old WT mice, the density of myelinated axons is significantly reduced compared to 6-month-old mice, and the percentage of fibres with a degenerate profile is significantly increased (860-1300 axons per nerve were examined). Data were analysed by unpaired, two-tailed Student’s t-tests (n = 5 or 6; *** p < 0.001, **** p<0.0001). **C.** Micrographs and graphs showing that APP positive axonal swellings and Iba1-labelled microglia are increased in 24-month-old mice compared to their 6-month-old counterparts in the corpus callosum (cc) but not in the motor cortex (mCtx), Bregma 0.74 mm. Mac3 positive microglia were significantly increased in both cc and mCtx in aged compared to younger mice. Data were analysed by unpaired, two-tailed Student’s t-tests for each region of interest (n=3; * p < 0.05, ** p<0.01, **** p<0.0001) **D.** Western blot showing that MCT1 is about 50% decreased in a “myelin-enriched” brain fraction from 24-month-old mice, compared to 6-month-old mice (top), using fast green (FG) as an internal standard. Data were analysed by an unpaired, two-tailed Student’s t-test (n = 5; ** p < 0.01).

When brain sections were stained for amyloid precursor protein (APP) as a marker of axonal transport impairment, we found a significant increase in APP positive swellings in the corpus callosum of 24-month-old mice (Fig. 7C), but interestingly not in the cortex. These changes were accompanied by increased densities of Iba1 and Mac3 positive microglia/macrophages. Notably, by western blotting, steady-state level of MCT1 in myelin enriched fractions was approximately 50% reduced in the 24-month-old mice compared to 6-month-old controls (Fig. 7D). Taken together, these data suggest that aging-associated changes of myelin integrity impairs normal transport processes in the myelinic channel system, having a profound impact on MCT1 levels, and on oligodendroglial metabolic support of axonal integrity.

## Discussion

The physiological function of compacted myelin in axonal insulation and impulse conduction is well established, but a distinct role for “non-compacted” myelin has been obscure. What is recognized by EM as non-compacted myelin was identified already at the light microscopic level by Rio de Hortega et al (1919), but suspected an experimental artifact (Edgar et al., 2021). Before the application of high-pressure freezing and freeze-substitution techniques to reduce EM dehydration artifacts (Edgar et al., 2021; Möbius et al., 2016), the relative volume of inner and outer tongues was grossly underestimated. While microtubules and membranous structures were observed in non-compacted myelin (Edgar and Griffiths, 2013), single EM cross-sections failed to show whether these are stationary remnants of development, myelin turnover or part of a vital transport system, analogous to axonal transport, between the oligodendrocyte soma and the innermost cytoplasmic myelin layer facing the axon. Here, by combining EM, mouse genetics, live imaging of peroxisomes and pharmacological interventions, we have established the concept of motor-driven, microtubule-dependent organelle transport in established myelin sheaths, being critical for long-term axonal integrity. Moreover, *in vitro* findings demonstrating that neuronal electrical activity affects myelinic transport suggests that myelinated neurons themselves can control, at least in part, aspects of oligodendrocyte behaviour.

Our interest in these phenomena was triggered by the observation that oligodendrocytes and axons are metabolically coupled (Fünfschilling et al., 2012) through MCT1, a monocarboxylate transporter localized at the glial axonal interface (Lee et al., 2012). While it is unclear currently whether myelin turnover in adult mice rests entirely on the incorporation of new membranes at the innermost myelin layer (Meschkat et al., 2020), i.e. reflecting developmental myelination (Snaidero et al., 2014), the necessary turnover of MCT1 and its functional incorporation at the periaxonal myelin membrane would require vesicular transport to the inner tongue. We thus hypothesized that MCT1 levels in mature myelin depend on the integrity of the intra-myelinic transport route. Our data support this suggestion by demonstrating a severe depletion of MCT1 and increased frequencies of organelles in myelin from CNP-deficient mice in which the myelinic channel system is structurally impaired (Snaidero et al., 2017); findings replicated in aged animals. In particular, the ‘trapping’ of myelinic organelles is reminiscent of organelle accumulations in axons upon perturbation of axonal transport (reviewed in Stassart et al., 2018).

To model and visualize transport of such nuclear-encoded membrane-bound cargoes in the myelinic channel system, we chose transgenic *Cnp-EOS^SKL^*-mice that harbour fluorescent peroxisomes in oligodendrocytes (Richert et al., 2014). These single membrane organelles are abundant in myelinating glia and while used here as a surrogate for myelin organelles more generally, it is noteworthy that oligodendroglial peroxisomes are essential for long-term axonal integrity (Kassmann et al., 2007).

The existence of a transport system operating in mature fibres is plausible because the myelinic channel system is developmentally derived from subcellular compartments of oligodendrocytes in which transport processes contribute directly to myelin outgrowth (Herbert et al., 2017; Iyer et al., 2024; Lyons et al., 2009; Nawaz et al., 2015; Snaidero et al., 2014; Yergert et al., 2021). Analogous to neurons with growth cones, oligodendrocytes in monolayer culture have microtubules within growing cellular processes (Barry et al., 1996; Song et al., 1999) with the notable difference that these processes remain short and deliver new membranes to a flat bilayer structure that, *in vivo*, grows spirally around axons (Snaidero et al., 2014). Indeed, *in vivo*, normal myelin sheath growth requires nucleation of microtubules, that is mediated by tubulin polymerization promoting protein (TPPP) (Fu et al., 2019; Lehotzky et al., 2010; Tokési et al., 2010). Further, mRNAs encoding myelin proteins (See Supplementary Fig.1) and presumably the oligodendroglial translation machinery, are transported on microtubules, as demonstrated in oligodendrocyte monolayer cultures (Brophy et al., 1993; Carson et al., 1998; Colman et al., 1982; Holz et al., 1996; Kursula et al., 2001; Quraishe et al., 2016; Trapp et al., 1987).

Considering our findings reported here, the importance of transport processes for developmental myelination and myelin maintenance is illustrated by the spontaneous *taiep* mutant rat, which harbours a point mutation in *Tubb4a*, encoding β-tubulin IV. This increases the ratio of “minus-end-distal” to “plus-end-distal” microtubules in the fine processes of developing oligodendrocytes (Song et al., 1999), which leads to an abnormal distribution of various myelin gene products (O’Connor et al., 2000) and hypomyelination with subsequent demyelination (Lunn et al., 1997).

Remarkably, the *Tubb4a* null mutation had no overt effect on myelination, indicating compensatory functions by other tubulins, and a dominant-negative effect of the *taiep* mutation. Nonetheless, the oligodendrocyte-specific expression of this tubulin isoform in the CNS (Schaeren-Wiemers et al., 1995) suggests that its known function as a tubulin stabilizer (Renthal et al., 1993) is important for myelinating oligodendrocytes.

A growing body of evidence has shown that neuronal electrical activity can stimulate developmental myelination (reviewed in Taylor and Monje, 2023). Less is known about axon-myelin signalling in the mature fibre, due in part to the inaccessibility of the glial-axonal junction on the inside of the myelin sheath. Our finding that the mobility of peroxisomes in myelinic channels is higher in electrically active networks, which, being a rapid response, suggests it involves direct axon-myelinic signalling. While mechanistic studies await the analysis of corresponding mouse mutants, we note that mature oligodendrocytes respond to axonal spiking activity (Yamazaki et al., 2010), at least in part through an NMDA receptor-mediated calcium rise in the myelin sheath (Micu et al., 2018) that also activates the intracellular transport of the glucose transporter GLUT1 in oligodendrocytes (Saab et al., 2016). Moreover, potassium transients in the periaxonal space of fast spiking fibres induce Ca^2+^ signals that stimulate glycolysis in oligodendrocytes (Looser et al., 2024).

The novelty of our findings is not the “myelinic channel system” as such, because we and others already hypothesized a “continuity” of non-compacted myelin as a cytosol-filled rim that flanks compacted myelin, with obvious reference to its developmental origin (Edgar et al., 2009, 2021; Hirano and Dembitzer, 1967; Nave, 2010b; Snaidero et al., 2014) and as a route for diffusing energy-rich metabolites for axonal support (Saab and Nave, 2017). Moreover, the same cytosolic spaces were previously visualized *ex vivo* by dye injections in both the CNS and PNS (Balice-Gordon et al., 1998; Butt and Ransom, 1989; Velumian et al., 2011). Rather, we provide the first experimental evidence that this system serves active transport processes, utilizing motor proteins and tubular tracts, exemplified by peroxisome movement. While this first description of “myelinic transport” (in analogy to “axonal transport”) is necessarily incomplete with respect to the nature of the motor proteins and tubulin isoforms involved, it already helps to explain aspects of the obscure secondary axonal degeneration of myelin-specific disorders. Adult CNP-deficient mice, modelling a rare human neurological disease (Al-Abdi et al., 2020), lack integrity of this myelinic channel system, as CNP normally prevents (MBP-dependent) myelin hyper-compaction and local “channel collapse” (Snaidero et al., 2017). Consequently, nuclear encoded MCT1 of mature oligodendrocytes probably fails in the motor-driven transport via the myelinic channel system to replenish supplies at the inner tongue of myelin as suggested by western blotting of a myelin-enriched tissue fraction. Loss of MCT1 in turn predicts poor metabolic interaction with axons (Lee et al., 2012), progressive axonal degeneration and the previously described premature death of the animal (Lappe-Siefke et al., 2003).

Clinically more relevant are myelin diseases, such as multiple sclerosis (MS), where immune-mediated attack on the myelin sheath causes progressive axon loss. We recently found that in human MS and its mouse models, a major contributor to secondary axonal degeneration is myelin injury (Schäffner et al., 2023). We hypothesise this has the same effect on MCT1 turnover as CNP deficiency, explaining the delayed axonal degeneration occurring months after the primary lesion (Schäffner et al., 2023).

The known neuroprotective function of oligodendrocytes involves the transfer of materials from the oligodendrocyte to the axon (Chamberlain et al., 2021; Frühbeis et al., 2020; Fünfschilling et al., 2012; Lee et al., 2012; Mukherjee et al., 2020). Thus, the oligodendrocyte acts locally, at the level of each myelinated internode, to support the axon (Edgar and Garbern, 2004; Edgar et al., 2004), which, being encased in lipid-rich myelin, is physically isolated from the extracellular milieu (Nave, 2010a). At the inner tongue of normal-appearing fibres in wild-type animals, we observed multivesicular bodies, as reported previously (Frühbeis et al., 2020) and fusion profiles (**Supplementary** Fig.1G) that could empty exosomes and soluble content, respectively, into the periaxonal space for uptake by the axon.

The brain’s white matter, which contains densely packed, parallel arrays of myelinated axons, is vulnerable during ageing in humans, and small diameter myelinated fibres are particularly vulnerable (Marner et al., 2003; Tang et al., 1997). If one assumes axon energy insufficiency is a pathological driver, this seems at first counter intuitive because thin axons are predicted to require less energy than thicker ones (Perge et al., 2012). A possible explanation relates to the fact that oligodendrocytes that myelinate thin axons extend many more processes than those that myelinate thick axons (reviewed in Edgar et al., 2021). As oligodendrocytes in the ageing brain can develop inclusion-containing bulbous swellings (presumably sites of transport stasis) along their processes (Peters, 2002), it becomes apparent that thin myelinated fibres might be more susceptible due to a greater risk of transport impairment in oligodendrocytes that extend numerous processes to small diameter axons.

## Limitations of the study

Direct visualisation of organelle transport in myelin was conducted in an *in vitro* model of the myelinated CNS, in which compact myelin with an appropriate g-ratio is formed (Thomson et al., 2008), and in which multiple paranodal loops abut the axon at the paranode (Edgar et al 2021 and **Supplementary Videos 4 and 5**). Like the CNS *in vivo*, these myelinating cell cultures also contain astrocytes, microglia and oligodendrocyte progenitor cells. Nonetheless, our findings should be confirmed *in vivo*.

A second limitation is that we cannot directly correlate myelin peroxisomal transport with the firing rate of the same axon. However, neurons exhibited a range of firing frequencies, corresponding to the wide variability in the proportion of myelin peroxisomes that exhibited movements at the time of imaging. Notably, myelin peroxisome motility was not completely blocked in the presence of TTX, but never reached the mobility seen in cell cultures with intrinsic neuronal electrical activity. Additionally, the number of organelles quantifiable in the inner tongues of myelin *in vivo* is small (estimated at ∼2 per sheath) which limits our ability to quantify changes following electrophysiological stimulation of optic nerve. Finally, the observed loss of MCT1 in a ‘myelin-enriched’ spinal cord lysates by western botting provides no direct evidence that the protein is reduced at the inner lip of the myelin sheath.

## Acknowledgements

We are grateful to Professors Ueli Suter and Pierre Chambon, and Dr Daniel Metzger for gift of the PLP-Cre-ERT2 mice. Dr Katherine Kusch (Nave lab) and Professor Ori Peles, Weizmann Institute of Science, kindly provided antibodies to MCT1 and Caspr1, respectively. Professor Lukas Kapitein, Utrecht University generously provided the plasmid encoding StableMARK and mNeonGreen, the latter provided by Allele Biotechnology & Pharmaceuticals. TagRFP-T_A1aY1 (158751) was provided to Addgene by Professor Minhaj Sirajuddin, Institute for Stem Cell Science & Regenerative Medicine. EM processing and sectioning of conditional *Kif21*b and *Tubb4a* knockout mice was performed with outstanding technical assistance from Mrs Margaret Mullin in the Glasgow Imaging Facility; We are grateful to colleagues in Glasgow, i.e. Dr Paul Montague for performed genotyping; Dr Fatma Kok and Mr David Kerrigan in Dr Rhaidhri Carmody’s lab for providing Midi-preps; John Cole for checking calculations for peroxisome densities; also to Dr Michaela Schweizer and Dr Mary Muhia (Hamburg) for assisting in respect to the conditional *Kif21b* knockout mice. J.M.E. and E.R.B. were supported by grants from the UK MS Society (Grants 58 to JME; Grant 127 to JME and ERB). J.M.E. was also supported by funds from the Chief Scientist’s Office, Scotland and the University of Glasgow (SPRINT MND/MS PhD studentship) and the National Centre for the Replacement, Refinement and Reduction of Animals in Research (NC3Rs). K.A.N. was supported by the European Research Council (AdvGrant MyeliNANO), the Dr. Myriam and Sheldon Adelson Medical Foundation (AMRF) and the German Research Council (DFG-TRR274). SG was supported by the Deutsche Forschungsgemeinschaft GO 2463/1-1.

## Authors’ contributions

Conceptualization JME, KAN, SG, ERB.

Data curation, KJC, JME.

Formal Analysis KJC, SW, Y-H C, UG, MLA, YK, ERB, JME.

Funding acquisition, JME, KAN, SG, ERB.

Investigation KJC, SW, UG, MLA, YK, LP-F, ERB, LK, JME.

Methodology AB, TG, CLC, RSS, LK, REM

Project administration JME, KAN, SG

Resources MK, CK, HBW

Supervision JME, SG, ERB

Visualization, JME, KJC

Writing – original draft JME, KAN

Writing – review & editing SG, ERB, KJC, MK, CC, HW, ID, all authors

All authors approved the manuscript

## Materials and methods

### Mice

Conditional *Kif21b* knockout transgenic mice and ‘reporter’ mice harbouring for various combinations of tamoxifen-inducible PLPCreERT2 (Leone et al., 2003; Weber et al., 2001), the “stop-floxed” Cre reporter gene ß-actin-tdTomato (Madisen et al., 2010) and CNP-PexEOS2, were maintained in the Biological Services Facility at the University of Glasgow, UK, under project licence PPL P78DD6. All regulated procedures involving these animals were approved by the University of Glasgow Ethical Review Committee, licenced under the UK Animals (Scientific Procedures) Act of 1986, and conducted according to the ARRIVE guidelines. Mice were housed in standard plastic cages or within individually ventilated cages, with 1-4 littermates, and nesting domes. They were maintained with a 12-hour dark/light cycle at 19-23°C, relative humidity 55% (± 10 %) and provided food and water *ad libitum*. *Tubba4* conditional knockout mice were bred and housed in the animal facility of the Max Planck Institute for Multidisciplinary Sciences (MPI-NAT). They had *ad libitum* access to water and food in standard plastic cages with 1-5 littermates in a temperature-controlled room (approx. 21°C) under a 12-hour dark/light cycle. Animal experiments were performed in accordance with the German animal protection law (TierSchG) and approved by the Niedersächsisches Landesamt für Verbraucherschutz und Lebensmittelsicherheit (LAVES) under license 33.19-42502-04-17/2409, if subject to approval. Procedures were supervised by the animal welfare officer for the MPI-NAT, Göttingen, Germany. The animal facility is registered at the MPI-NAT is registered according to §11 Abs. 1 TierSchG. CD-1 mice were purchased from Charles River Laboratories (Margate, UK) and housed at the University of Nottingham with *ad libitum* access to food and water, temperature 22–23°C, and a 12-hour dark/light cycle.

Transgenic reporter mice: Male and female pairs, which together harboured one copy of each of three transgenes, were mated/time-mated to produce embryos/offspring harbouring combinations of transgenes encoding tamoxifen-inducible PLPCreERT2 {Leone, 2003 7898 /id; Weber, 2001 7163 /id}, the “stop-floxed” Cre reporter gene ß-actin-tdTomato {Madisen, 2010 10467 /id} and CNP-PexEOS2 {Richert, 2014 10701 /id}. Approximately one in eight embryos/offspring harboured all three transgenes leading to the expression of EOS2 +ve peroxisomes in tdTomato +ve oligodendrocytes.

*cKif21b* KO mice (Muhia et al., 2016): By intercrossing *Kif21b^lacZ/neo^*mutants with mice expressing FLIP recombinase (129S4/SvJaeSor-*Gt(ROSA)26Sor^tm1(FLP1)Dym^/J*; backcrossed into C57BL/6N for at least 10 generations) (Farley et al., 2000) the lacZ/neo cassette was excised *in vivo*, yielding the “conditional ready” heterozygous flox allele in mice carrying the *Kif21f* KIF21btm2a(KOMP)Wtsi allele (also termed *Kif21b^flox/+^*). *Tubb4a^flox/+^* mice were subsequently bred to homozygosity. To inactivate expression of *Kif21b* in oligodendrocytes, homozygous mice were crossed with mice expressing *Cre* under the control of the *Cnp* promotor (*Cnp^tm1(cre)Kan^*) (Lappe-Siefke et al., 2003) to generate *Cnp*^+/Cre^::*Kif21b^fl/fl^*; *Cnp*^+/Cre^::*Kif21b*^+/+^ or *Cnp*^+/Cre^::*Kif21b*^+/+^; or *Cnp*^+/+^::*Kif21b*^fl/fl^ offspring.

*Tubb4a* conditional knockout mice: Embryonic stem cell clones HEPDD0853_6_E02 and HEPD0853_6_B04 harboring the tubulin beta 4A class IVA *Tubb4a^tm1a(EUCOMM)Hmgu^*‘knockout-first’ allele (reporter-tagged with conditional potential) of the Tubb4a gene with exons 2 and 3 flanked by loxP sites were obtained from the European Mouse Mutant Cell Repository (EuMMCR, Munich, Germany) and microinjected into blastocysts by standard procedures. Highly chimeric animals were crossed to C57Bl/6N females yielding *Tubb4a^tm1a(EUCOMM)Hmgu^*mutant mice (also termed *Tubb4a^lacZ/neo^).* By intercrossing *Tubb4a^lacZ/neo^* mutants with mice expressing FLIP recombinase (129S4/SvJaeSor-*Gt(ROSA)26Sor^tm1(FLP1)Dym^/J*; backcrossed into C57BL/6N for at least 10 generations) (Farley et al., 2000) the lacZ/neo cassette was excised *in vivo*, yielding the “conditional ready” heterozygous flox allele in mice carrying the *Tubb4a^tm1c(EUCOMM)Wtsi^*allele (also termed *Tubb4a^flox/+^*). *Tubb4a^flox/+^*mice were subsequently bred to homozygosity. To inactivate expression of *Tubb4a* in oligodendrocytes, the loxP flanked exons 2 and 3 were excised *in vivo* upon the generation and appropriate intercrossing of *Tubb4a^flox//flox^*mice with phenotypically normal *Tubb4a^+/+::^CNP^Cre/+^* mice that expressed *Cre* under the control of the *Cnp* promotor (*Cnp^tm1(cre)Kan^*) (Lappe-Siefke et al., 2003).

For optic nerve electrophysiology, CD-1 male mice (weight 28–35 g, corresponding to 30– 45 days of age) were purchased from Charles River Laboratories (Margate, UK) and housed at the University of Nottingham with *ad libitum* access to food and water and temperature 22–23°C, on a 12-hour dark/light cycle. Experiments were approved by the University of Nottingham Animal Care and Ethics Committee and carried out in accordance with the Animals (Scientific Procedures) Act 1986 under appropriate authority of establishment, project and personal licences.

### Genotyping

Routine genotyping was performed by genomic PCR on ear clip DNA. Ear notches were each lysed in 100 µL 50 mM NaOH at 95°C for 90 minutes, then neutralized with 10 µL 1M Tris buffer (pH 5) followed by thorough vortexing.

*Tubb4a^+^ and Tubb4a^flox^* alleles were detected using sense primer P1 (5’-GAC AGT CAG CAA CTG AAG AAG TGG C-3’) together with antisense primer P2 (5’-CTG TTG TCA ATC TTG GCC ATG AGC C-3’) and P3 (5’-TGA ACT GAT GGC GAG CTC AGA CC -3’) amplifying a 546 bp product for the WT allele, a 748 bp product from the flox allele, and a 674 bp product from the recombined flox allele.

*Cnp^Cre^ and Cnp^wt^* alleles were identified by genomic PCR with primers P4 (5’-GCC TTC AAA CTG TCC ATC TC-3’), P5 (5’-CCC AGC CCT TTT ATTA CCA C-3’), and P6 (5’-CAT AGC CTG AAG AAC GAG A-3’) amplifying a 700 kb product from the wt allele and a 400 bp product from the Cre or with *Cnp1* Sense 5^/^-GCC TTC AAA CTG TCC ATC TC-3^/^, Cnp1N2 5^/^-GAT GGG GCT TAC TCT TGC-3^/^ and Puro3 5^/^-CATAGCCTGAAGAACGAGA-3^/^, amplifying a 1180 bp and 894 bp products for Cnp1^WT^ and Cnp1^Cre^, respectively.

Mice expressing tamoxifen-inducible tdTomato fluorescent protein and mEOS2-lablelled oligodendrocyte peroxisomes were genotyped using the following primers:

Tom Fw 5^/^ TAC GGC ATG GAC GAG CTG TAC AAG TAA-3^/^ and Tom Rv 5^/^ CAG GCG AGC AGC CAA GGA AA-3^/^, yielding an ∼500bp product.

EOS Fw 5^/^-CTT CTT ACA CAG GCC ACC ATG AGT GCG-3^/^ and EOS Rv 5^/^-GGA TCC TTA CTT AGT TAA AGC TTG GAT CGT-3^/^, yielding an ∼800bp product.

PLP1 Fw 5^/^ TGG ACA GCT GGG ACA AAG TAA GC and PLP Rv 5^/^ CGT TGC ATC GAC CGG TAA TGC AGG C-3^/^ yielding an ∼250 bp product.

*Kif21b*^+^ and *Kif21b*^flox^ alleles were identified by genomic PCR with primers Fw Kif 83 5^/^-GGA CAT GAT GGT GAT TTA GAG G-3^/^ and Rv Kif84 5^/^-AAA CCT GGG CAA AGG CAT AC-3^/^ yielding 825 bp (WT) and 970 bp (flox allele) products.

### Transmission electron microscopy of conventionally fixed tissue

Transcardiac perfusion of mice was performed under the effect of a lethal dose of Avertin or Dolethal. For transmission electron microscopy (TEM), mice were perfused with 4% paraformaldehyde (PFA), 2.5% glutaraldehyde in 0.1 M phosphate buffer, postfixed in 4% PFA overnight and subsequently transferred to 1% PFA (conditional *Tubb4a* knockout mice and controls) or with 4% PFA and 5% glutaraldehyde in cacodylate buffer, and postfixed in the same for 24 hours to three weeks (c*Kif21b* KO mice and controls). Optic nerves were subsequently dissected and processed for EM, as described (Edgar et al., 2020).

Following electrophysiology of CD1 mouse optic nerves, paired nerves (stimulated and non-stimulated) were immersed in 4% PFA and 5% glutaraldehyde in cacodylate buffer for at least one week and processed and sectioned as above. Morphometry and quantification of electron micrographs were performed using Fiji (Schindelin et al., 2012).

### Transmission electron microscopy of high-pressure frozen tissue

Tissue was collected and processed as described in (Möbius et al., 2016).

### White matter morphometry

Optic nerve cross sections taken mid-way between the retina and the extraorbital region of the nerve were imaged at 12000 x on a JEOL 1200 EX transmission electron microscope (c*Kif21b* and c*Tubba4* KO mouse optic nerves) or an EM912AB electron microscope (aged mice). Images of optic nerve cross section were captured randomly at 12,000x magnification using Olympus ITEM Software (12-20 images per nerve, each ∼74 mm^2^) or at 3150x (5 images per nerve, each 480 mm^2^) using the ImageSP software (TRS, SysProg, ImageSP, vers1.1.4.62), respectively. Images were opened in Fiji (Schindelin et al., 2012), and the relevant feature was counted using the Image J Cell Counter plug-in within and on west and south borders of an area of interest (AOI), by a blinded experimenter. The sum of the counts from all micrographs from a single nerve was converted to numbers per mm^2^. Some counts were converted to fibres per nerve after measuring the area of the optic nerve cross section from toluidine blue stained resin sections. For quantifying organelles in inner tongue and paranodal loops, dense organelles (Fig. 1) were identified, and the numbers of organelles was divided by the number of myelinated fibres to determine the number of organelles per fibre cross-section. To determine organelle diameter, the area of each organelle was measured in Fiji (Schindelin et al., 2012) by tracing the organelle bounds and the diameter calculated, assuming the area was circular. Organelles were assumed to be spherical, and the maximum diameter observed on optic nerve cross sections was assumed to represent the maximum diameter in the longitudinal direction (along the sheath).

### Tissue collection for light microscopy (frozen sections)

Mice were perfusion fixed under the effect of a lethal dose of Dolethal, with 0.9% saline followed rapidly by 4% PFA and immersed in 4% PFA for a minimum of 24 hours before the optic nerve and spinal cord were dissected. Dissected tissues were transferred into 20% sucrose in PBS until they sank. Tissues were frozen in OCT (Sakura Finitek, Torrance, CA, USA) in liquid nitrogen-chilled isopentane. Ten µm thick sections were cut on a cryostat (Bright Instruments) and mounted on Colorfrost Plus charged slides (Thermo Fisher Scientific). Tissue sections were permeabilised in methanol for 10 minutes at -20°C and immunostained with rabbit anti-Caspr1 (kindly gifted by Professor Ori Peles) by incubation overnight at 4°C in blocking buffer (10% goat serum, 1% bovine serum albumin (BSA), 0.1% Tween, 0.9 % NaCl in PBS). Bound primary antibodies were detected using Alexa 647 goat anti-rabbit IgG (1:1000; Invitrogen).

### Tissue collection for light microscopy (paraffin sections)

Mice were anesthetized with a lethal dose of avertin and perfused with Hanks balanced salt solution followed with 4% PFA in 0.1 M phosphate buffer. The brains were dissected, post-fixed overnight in 4% PFA, paraffinized using a Microm STP 120 Spin Tissue Processor (Thermo Scientific) and subsequently embedded in paraffin blocks on a HistoStar embedding workstation (Epredia).

### Immunohistochemistry and quantification on paraffin sections

Five[μm brain sections in paraffin were cut on a RM2155 (Leica) rotary microtome and mounted on Histobond adhesive microscope slides. Slices were deparaffinized at 60°C and rehydrated using xylene and a descending ethanol series. For heat-induced epitope retrieval, sections were treated in sodium citrate buffer (0.01 M, pH 6.0) in a 400-Watt microwave for 10 minutes, followed by a 20-minute cooldown period. For immunohistochemical antibody detection using a chromagen, endogenous peroxidase activity was blocked with 3% hydrogen peroxide. Subsequently, sections were blocked for 1 hour at RT in 20% goat serum in PBS and incubated with primary antibodies (anti-MAC3, anti-Iba1, anti-APP) at 4°C overnight. The LSAB2 system-HRP kit (DAKO) was use and the HRP substrate 3,30-Diaminobenzidine (DAB) was applied using the DAB Zytomed kit (Zytomed Systems GmbH). Nuclei were labeled by Hematoxylin stain. Imaging was performed using a Zeiss Axio Imager Z1 bright-field light microscope equipped with a Zeiss AxioCam MRc camera using ZEN 2012 blue edition software. Images of anterior corpus callosum and motor cortex (Bregma 0.74 mm) were color deconvoluted using the “Color Deconvolution” (1.3) plugin in Fiji/ImageJ. Iba1^+^ and MAC3^+^ areas were quantified using thresholding, and APP spheroids were quantified manually using the “Cell counter” plugin from Fiji/ImageJ. All data were normalized to the respective quantified areas.

### Single molecule FISH

Candidate smFISH probe sequences recognising mouse *Mbp* (NM_001025251) or *Mobp* (NM_001039365) were generated using Stellaris Probe Designer version 4.2 (https://www.biosearchtech.com/stellaris-designer) with the following parameters: organism, mouse; masking level, 5; oligo length, 18 nt; minimum spacing length, 3 nt. BLAST was used to remove potential off-target oligonucleotide sequences (e-value cutoff of 1e-2). The 3’ ends of oligonucleotides were labelled with Alexa Flour (Gaspar et al., 2017) normalised to 25 µM. All probe sets used had a degree of labelling > 0.99. Probe sequences are provided in **Supplementary Table 2**.

smFISH was conducted according to (Titlow et al., 2018) with minor modifications. At day *in-vitro* 28, myelinating cells cultures, adhered to #1 glass coverslips, were fixed in 4% paraformaldehyde for 10 minutes. Cells were washed twice in PBS and stored in 70% ethanol at 4° C until use. Cells were rehydrated in successive incubations (35%, 17.5% and 0%) ethanol 2’ SSC, for 5 minutes each RT. Pre-hybridization, cells were washed in pre-warmed wash solution (2× SSC, 10% formamide, 0.1% Tween-20) twice for 20 min each at 37°C. Hybridisation was carried out in 2× SSC, 10% formamide, 10% dextran sulphate, 0.1% Tween-20, containing 250 nM smFISH probes overnight at 37°C. Cells were then washed in pre-warmed wash solution for 20 minutes and stored in PBS. Prior to immunocytochemistry, RNASecure (Invitrogen) was added to Odyssey blocking solution (Li-Cor), and applied to cells for 1 hour RT. Cells were then incubated in primary antibody solution (anti-CNPase; Abcam) in Li-Cor blocking buffer overnight at 4°C. Cells were washed in PBS and incubated in Alexa flour 568 anti-mouse secondary antibody in Li-Cor blocking buffer (Invitrogen) for 1 hour RT. Cells were washed in PBS and incubated in TrueBlack Plus (Biotium) to reduce autofluorescence, according to the manufacturer’s instructions, and were then rinsed and mounted with Mowiol with DAPI.

Cells were imaged using an Olympus SpinSR (CSU-W1) spinning disk confocal microscope with a Prime BSI sCMOS camera. 100x oil (1.45NA, UPLXAPO100XO). Image voxel sizes were 0.067 × 0.067 × 0.2 µm (x:y:z).

### Quantitative RT-PCR

RNA was extracted from brain lysates and purified myelin-enriched fractions using the miRNeasy Mini Kit (Qiagen) and the concentration was increased using Co-precipitant pink (Bioline), if necessary. The purity and concentration of RNA was evaluated using a NanoDrop spectrophotometer and RNA Nano (Agilent). cDNA was synthesized using Superscript III (Invitrogen, Carlsbad, CA, USA) according to the manufacturers’ instructions. Quantitative RT-PCR (qRT-PCR) was performed in triplicates with the GoTaq qPCR Master Mix (Promega) on a LightCycler 480 Instrument (Roche Diagnostics). Expression values were normalised to the mean of the house-keeper genes *Atp1a1*, *Hprt* and *Rplp0*. Relative expression changes were analysed using the 2ΔΔC(T) method, normalised to mean expression in P20 wildtype myelin-enriched fractions (set to 1). All primers were designed with Universal Probe Library from Roche Applied Systems (https://www.roche-applied-science.com) and validated with NIH PrimerBlast. All primers were intron-spanning. Primers were specific for *SCL16a1/Mct1* (forward 5’-GGATATCATCTATAATGTTGGCTGTC-3’, reverse 5’-GCTGCCGTATTTATTCACCAA-3’; Atp1a1 (Forward 5’-GGCCTTGGAGAACGTGTG-3’, Reverse 5’-TCGGGAAACTGTTCGTCAG-3’); Hprt (Forward 5’-CCTGGTTCATCATCGCTAATC-3’, Reverse 5’-TCCTCCTCAGACCGCTTTT-3’) and Rplp0 (Forward 5’-GATGCCCAGGGAAGACAG-3’, Reverse 5’-ACAATGAAGCATTTTGGATAATCA-3’).

### Tissue collection and western blotting of myelin-enriched fraction and optic nerves

Male c57Bl6/N wildtype mice and CNP null mutant mice (Cnp^tm1(cre)Kan^; Lappe-Siefke et al., 2003) from breeding colonies at the Max Planck Institute of Multidisciplinary Sciences and male and female *Cnp1*^+/+^::*Kif21b*^flx/flx^, *Cnp1*^+/Cre^::*Kif21b*^+/+^ or *Kif21b* ^+/flx^ and *Cnp1*^+/Cre^::*Kif21b^flx/flx^* from breeding colonies at the University of Glasgow were used. A light-weight membrane fraction enriched for myelin was purified from mouse brain hemispheres or spinal cords, as described (Erwig et al., 2019). Optic nerves were homogenised in ROLB Buffer, pH 7.4 (10 mM HEPES, 50 mM NaF, 25 mM Na₄P₂O₇, 80 mM KH_2_PO_4_, 5 mM EDTA and 1% SDS with protease inhibitor) using a plastic pestle and mortar then centrifuged at 16,000g for 10 minutes, 4°C and the soluble fraction was aspirated. Protein concentration was determined using the DC protein assay (BioRad) or a BCA Protein Assay Kit (Thermo Fisher) according to the respective manufacturer’s instructions. Immunoblotting of brain ‘myelin’ was performed as described (Späte et al., 2024). Spinal cord ‘myelin’ and optic nerve samples were resolved on 4-12% Bis-Tris gel at 200 V and transferred onto PVDF membrane using either semi-dry method (Trans-blot; BioRad) or a wet gel transfer (Mini Trans-blot; BioRad). Semi-dry transfers were run at 1.3 mA/cm^2^ for 1 hour ± 15 minutes and wet gel transfers were run overnight at 35 V or for 1 hour at 100 V. Membranes were blocked with 5% milk in Tris buffered saline with 0.01% Tween 20 (TBS-T) then incubated in primary antibody in blocking buffer overnight at 4°C. Primary antibodies were specific for SCL16a1/MCT1 (1:50-1:1000, Stumpf et al., 2019), ATP1a1 (1:2000; Abcam) Kif21b (1:50; HPA027274, Sigma; adsorbed against *Kif21b* knockout brain tissue), CNP (1:2000; ab6319, Abcam), MOG (1:2000; gifted by Professor Christopher Linington), and β-Actin (1:5000; A1978, Sigma). Membranes were thoroughly washed (1X TBS/T) and binding was detected using appropriate peroxidase conjugated secondary antibody (Anti-mouse/rabbit IgG HRP-linked, Cell Signalling Technologies), wash was repeated then incubated for 1 minute in WesternBright^TM^ ECL mix (K-12045-D20, Advansta). Immunoblots were scanned using the Intas ChemiCam system or using the Li-Cor C-DiGit blot scanner.

### Myelinating cell cultures

Pregnant ‘reporter’ mice were killed on embryonic day 13 (E13; day of plug being E0) by cervical dislocation followed by exsanguination or CO_2_ exposure followed by cervical dislocation. Following removal of the uterine horns, isolation of the embryos and spinal cord removal, all embryonic cords were pooled and used to generate a single cell suspension for culture, as described (Bijland et al., 2019) resulting in a population of cells with a mixed genotype. Myelinating cultures were maintained for up to 50 days *in vitro* (DIV), in a cell culture incubator at 37°C in humidified 5% CO_2_. One µM (final) 4-hydroxy tamoxifen was administered for 1 hour on DIV 21 ± 3 days to induce tdTomato expression in a subpopulation of oligodendrocytes.

### Neuronal cell cultures

These were established as for myelinating cell cultures (Bijland et al., 2019), except that the dissociated spinal cord cells were plated in 80 µL serum-containing plating medium (50% DMEM, 25 % HBSS and 25 % horse serum) at 80,0000 cells per well in Seahorse XFp Miniplates (Agilent, 103025-100). After two hours, the serum-containing media was removed and replaced with 125 µL neurobasal medium (Gibco, A2477501) supplemented with B-27 (Gibco, 17504044), 10 µM 5-Fluoro-2^/^-deoxyuridine (FDU; Sigma-Aldrich, 343333), 10 ng nerve growth factor (NGF; Sigma-Aldrich, N6009), 10 mM glucose (Agilent, 103577-100), 0.5 mM glutamine (Agilent, 103579-100), 0.227 mM pyruvate (Agilent, 103578-100). The cells were maintained until DIV 15 ± 1 day and media was changed every 2 days.

### Seahorse neuromodulator assay on neuronal culture

The cartridges for the Seahorse XF Bioanalyser (Agilent) were prepared according to the manufacturer’s instructions. On the day of the assay, neurobasal media was changed to 180 µL of the Seahorse assay medium (Agilent, 103680-100, supplemented with 2 mM glutamine, 1 mM pyruvate and 10 mM glucose) and equilibrated in a non-CO_2_ incubator for 1 hour at 37°C. The assay was as follows: 3 cycles of baseline measurements were followed by injection of 100 μM PTX and two measurement cycles, followed by injection of 1 μM TTX and 2 measurements cycles. The total duration for the assay was 40 minutes and oxygen consumption rate (OCR) and extracellular acidification rate (ECAR) were recorded. The OCR and ECAR values for each phase were averaged and plotted over time.

### Treatment of myelinating cultures with nocodazole

On DIV 27 ± 3 days, myelinating cultures were treated for 3 hours at 37°C in a humidified 5% CO_2_ incubator with nocodazole (final concentration 20 μM; 1.2 μL of 5 mg/mL in DMSO was added per 1 mL media; SML1665; Sigma-Aldrich).

### Treatment of cultures with neuromodulators

On DIV 29 ± 5 days, myelinating cultures were treated for ≥ 5 minutes with either PTX (final concentration 100 μM; 1 μL stock solution in DMSO in 1 mL media; P1675; Sigma-Aldrich), or TTX (final concentration 1 µM; 1 µL stock solution in water in 1 mL media; 1078; Tocris, Abingdon, UK) A DMSO-only control was conducted for comparison with PTX.

### Live imaging of cell cultures

For live imaging, myelinating cultures were plated on custom-made imaging dishes, prepared from 35 mm diameter Petri-dishes (353001; Falcon), with three x 8 mm diameter circular holes in the base and a 25 mm diameter glass coverslip (630-2213; VWR) glued underneath (Bijland et al., 2019). Live tdTomato rendered oligodendrocytes with mEOS2-lablled peroxisomes were imaged using an inverted Zeiss Observer Z1 Spinning Disc confocal microscope, equipped with a Yokogawa CSU-X1 filter wheel and spinning disc unit, a Photometrics Evolve 512 delta EM-CCD camera, with 488 nm (300-600 ms exposure, 4-9 % power) and 568 nm (200-400 ms exposure, 4-6% power) laser lines and a 63x/1.4 Oil Pln Apo objective and Zen software (2012, Blue Edition). Alternatively, an inverted Leica DMi8 widefield microscope, with a Hamamatsu ORCA-Flash4.0 V3 Digital camera C13440-20C, HC PL APO CS2 63x/1.40 Oil objective, and LAS-X software (version 3.7.1) or an inverted Zeiss Observer Z1 epi-fluorescence microscope equipped with an AxioCam MRc3, ×0.63 Camera Adaptor, ZEN 2012 blue edition software using a Zeiss plan-apochromat 63 × 1.4 oil objective, were used. Cells were maintained at 37°C on a heated stage insert and gassed with humidified 5% CO_2_ to maintain physiological pH, which was monitored on the basis of the colour of the phenol red-containing cell culture media. To record peroxisome movement, frames were captured every 10 seconds over a 10 minute period, in the 568 (tdTomato) and 488 (mEOS2) channels.

### Quantification of peroxisome motility

All quantification was performed blinded using time-lapse images of tdTomato +ve, myelinating oligodendrocytes containing mEOS2-labelled peroxisomes. Numbers of peroxisomes in oligodendrocyte processes or sheaths were counted manually using the multi-point tool in Fiji (Schindelin et al., 2012). To calculate the percentage of motile peroxisomes in oligodendrocyte processes or sheath, the total number of peroxisomes and the number of motile peroxisomes were recorded. Peroxisomes that moved ≥3 µm were considered ‘motile’ and cells were excluded if there were <5 myelin peroxisomes observable in the time-lapse image. To measure the distances travelled by the motile peroxisomes, the 488 (mEOS2) channel of the time-lapse images was analysed using the Simple Neurite Tracer plugin (https://imagej.net/plugins/snt/) with Fiji. Time lapse frames were converted into an AVI file, and the point tool in Simple Neurite Tracer was used ‘mark’ the position of the peroxisome at each frame, and distances travelled between frames, including bidirectional movements, were summed. Average speed of each peroxisome was calculated by dividing the total distance travelled (µm) by time (seconds). Motile speed was calculated by dividing the total distance travelled by the number of frames in which the peroxisome was motile.

For motile peroxisomes in processes (between the soma and the sheaths), the direction of movement in relation to the cell body was also recorded, and the net distance travelled in each direction (anterograde or retrograde) was quantified. First, the length of each process was measured using the Simple Neurite Tracer plugin. Second, as process lengths varied, and hence, impacted distance available, the distance travelled by each peroxisome (measured as above) was normalised to distance per 100 µm process. Third, to calculate the net movement in the anterograde or retrograde direction per process, the normalised distance travelled per peroxisome was multiplied by 1/total number of peroxisomes per process (including stationery ones) and then the distances in either direction were summed. If there were no motile peroxisomes in a process, zero was recorded for both anterograde and retrograde movement. If there was movement in only one direction, zero was recorded for the opposite direction. For example, for a process with 5 peroxisomes, of which 1 moved 50 μm per 100 μm in the anterograde direction, the net distances were defined as anterograde: 1/5 x 50 μm = 10 μm per 100 μm and retrograde: 0 μm per 100 μm.

### Culture on nanofibres

Spinal cords were dissected from postnatal day 3-5 (P3-5) ‘reporter’ mice, digested using trypsin (0.125%) in HBSS minus Ca^2+^ and Mg^2+^, and dissociated into a single cell suspension by trituration through 21 and 23 G needles. Only glial cells survive this process, eliminating almost all neurons from the cell culture. Cells were plated at 50,000 cells per well on 4 μm diameter Poly-L-Lactic Acid (PLLA) electrospun fibers in 12-well cell crown scaffolds (The Electrospinning Company) in serum-containing plating medium (see Myelinating Cultures). The following day, the plating medium was removed and the glial cells were maintained in Sato’s medium (DMEM, glucose [25 mM], pyruvate [1 mM], penicillin/streptomycin [1% v/v], apo-transferrin [5 μg/ml], sodium selenite [5 ng/ml], insulin [5 μg/ml], putrescine [16 μg/ml], progesterone [7.3 ng/ml], T3 [0.5 nM] and horse serum [1% v/v]), with media replenished 50:50 every 7 days.

### Immunocytochemistry

Cell cultures were fixed in 4% PFA at RT for 10 minutes, or by the addition of 8% PFA (37°C) to cell culture media (1:1 v:v), then permeabilised for 10 minutes in either 0.5% triton X at RT or in absolute methanol at -20°C. Following washes in PBS, blocking buffer (10% goat serum, 1% bovine serum albumin (BSA), 0.1% Tween, 0.45 g NaCl in 50 mL PBS) was applied for 1 hour at RT and then cells were incubated overnight in primary antibody at 4°C, in 10% goat serum in PBS or blocking buffer. After thorough washing, secondary antibodies were applied at 1 in 1000 for ∼1 hour at RT. Finally, cells were washed in PBS and rinsed in water, then coverslips were mounted on glass slides in Mowiol containing DAPI (2 µg/ml; Invitrogen). For immunocytochemistry of microtubules, cells were first incubated for 1 minute with pre-warmed extraction buffer (0.3% Triton X and 0.1% glutaraldehyde in BRB80 buffer (BRB80 buffer: 80 mM Pipes, 1mM EGTA and 4 mM MgCl_2_, pH 6.8) (Jansen et al., 2021) then fixed for 10 minutes in 4% PFA and permeabilised for 10 minutes in 0.25% Triton-X before blocking for 1 hour in 3% BSA/PBS at RT. Cells were next incubated for 1 hour with primary antibody at RT, diluted in 3% BSA/PBS, washed in PBS, then incubated for 1 hour at RT with secondary antibodies diluted in 3% BSA/PBS. Coverslips were washed and mounted as described above. Primary antibodies were; mouse IgG2a anti-β tubulin III (1:200; T8578; Sigma), mouse IgG1 anti-β tubulin IV (1 in 400; ab11315; Abcam), rat anti-MBP (1 in 800; ab7349; Abcam), mouse IgG1 anti-CNP (1 in 800; ab6319; Abcam), rabbit anti-Caspr1 (1:1000; kindly provided by Professor E Peles), mouse IgG2a anti-Ermin (1:500; MABN323; Merck); mouse IgG3 anti-tyrosinated tubulin (1:300; T9028; Merck); mouse IgG2b anti-acetylated tubulin (1:300; T7451; Merck); rabbit anti-detyrosinated tubulin (1:300; ab48389; Abcam). Bound primary antibodies were detected using Alexa 488, 568 or 647 goat anti-mouse IgG, IgG1, IgG2a, goat anti-rabbit IgG or goat anti-rat IgG (1:1000; Invitrogen).

To stain cells cultured on nanofibres, the nanofibre crown was transferred between wells of a 12-well dish for permeabilisation of cells, antibody incubations and washing, as above. The nanofiber crowns were placed on glass slides and the fibres were sandwiched between the slide and a 13 mm glass coverslip, using Mowiol with DAPI. The crown was removed 2 days later, after the Mowiol had hardened, as described (Bechler, 2019).

### Transfection with sensors of microtubule post translational modifications

Myelinating cultures were incubated with DharmaFECT 1 transfectant reagent (T-2001-01; Horizon) according to manufacturer’s instructions and with 1.125 µg/mL tyrosinated tubulin sensor TagRFP-T AlaY1 (158751; Addgene) (Kesarwani et al., 2020) or Stable Microtubule-Associated Rigor-Kinesin (StableMARK) (Jansen et al., 2023) for 9 hours. Cells were immediately fixed in 4% PFA/PBS and immunostained as described in ‘Immunocytochemistry’.

### Fluorescence Microscopy

Fixed cell cultures and tissue sections were generally imaged with an inverted Zeiss Observer Z1 epi-fluorescence microscope equipped with an AxioCam MRc3, x0.63 Camera Adaptor, ZEN 2012 blue edition software using Zeiss plan-apochromat 63 x 1.4 oil objective. High resolution images were taken using a Zeiss LSM 880 confocal microscope or Zeiss Z.1 AxioImager with ApoTome attachment, using Zen software and Plan-Neofluar 40x/1.30 oil objective or Plan-Neofluar 63x/1.30 oil objectives. Cells on nanofibres were imaged on an Olympus IX70 microscope, QIcam Fast digital camera (QImaging Surrey, BC, Canada) and Image-Pro Plus 6 software (Media Cybernetics, Silver Spring, MD).

### Quantifying peroxisomes in paranodes

Myelinating cultures were treated with neuromodulators, TTX or PTX (see Treatment of Cultures with Neuromodulators) for 5 minutes, then immediately fixed in 4% PFA/PBS (10 minutes) and either immunostained with anti-CNP (see Immunocytochemistry) and mounted on glass slides in Mowiol or mounted in Mowiol without immunostaining when expressing tdTomato. Following blinding, coverslips were scanned in the green (mEOS2-labelled peroxisomes) and red (oligodendrocytes stained with anti-CNP or labelled with tdTomato) channels, and oligodendrocytes with mEOS2-peroxisomes in the soma were selected at random. A z-stack (0.32 μm steps) was captured of paranodal regions using a Zeiss Observer Z1 epi-fluorescence microscope. Numbers of peroxisomes in each paranode were counted manually in Fiji or in Zen 2012 (Blue edition) in a minimum of 10 paranodes per treatment.

### Electrophysiology of optic nerve

Following decapitation, the optic nerves were cut behind the eye orbit then exposed by lifting the cerebral hemispheres rostro-caudally. They were cut at optic chiasm (5-7 mm length) and gently freed from their dural sheaths then placed in an interface perfusion chamber (Medical Systems, Greenvale, NY, U.S.A.), as described (Stys et al., 1991). Optic nerves were maintained at 37°C using a TC-202A bipolar temperature controller (Digitimer, Welwyn Garden City, UK) and perfused with artificial cerebrospinal fluid (aCSF) that contained (in mM): 153 Na^+^, 3 K^+^, 2 Mg^2+^, 2 Ca^2+^, 143 Cl^-^, 26 HCO_3_, 1.25 HPO ^2-^, and 10 glucose. The aCSF was bubbled with a gas mixture (95% O_2_: 5% CO_2_) to maintain pH at 7.45. Tissue was oxygenated by a humidified gas mixture (95% O_2_: 5% CO_2_) that flowed over its surface. Nerves were allowed to equilibrate for 30 minutes then a stimulating suction electrode back-filled with aCSF was attached to the proximal end of one nerve. A recording electrode filled with aCSF was attached to the distal end of the same nerve to record the compound action potential (CAP), which was evoked by a 125% supramaximal stimulus (30 µs in duration: a Grass S88 stimulator connected to two SIU5 isolation units in parallel capable of delivering two independently controllable pulses). Nerves were stimulated at 7 or 50 Hz for 20 minutes. CAPs were recorded from a suction electrode attached to the distal end of the nerve connected to an SR560 low noise preamplifier (Stanford Research Systems, Sunnyvale, CA 94089, USA): the signal was amplified up to 1000x and filtered at 30 kHz. The differential output was fed into a HumBug Noise Eliminator (Digitimer Ltd, Welwyn Garden City, UK) to remove mains frequency noise (50 Hz). The signal was acquired via an Axon Digidata 1550B using Clampex 11.3 (Molecular Devices Ltd, Wokingham, Berkshire, UK). The control nerve remained untouched on the perfusion chamber, and when the stimulation protocol was completed, both nerves were transferred rapidly to fixative as described in ‘Tissue collection and fixation for electron microscopy’.

### Electrophysiology of myelinating cultures

Myelinating cultures were transported between sites at RT in a sealed box containing humidified 5% CO_2_ (maximum journey time 1.5 hours) and allowed to equilibrate at least 1 hour in cell culture incubator (37°C, 5% CO_2_). One glass coverslip at a time was removed from the cell culture media dish and placed in a clean 35 mm diameter Petri dish containing pre-warmed extracellular solution, then viewed using an Olympus IX71 inverted microscope with epifluorescence. Whole cell voltage and current clamp recordings were performed on DIV 22-29 cultures using an Axopatch 200B amplifier with a National Instruments NIDAQ-MX digital acquisition system and results acquired and analysed with WinEDR V3.9.1 software. Experiments were performed at 35-37°C in atmospheric air using an extracellular solution containing the following ion concentrations (in mM) to mimic the potassium concentration of the cell culture media and concentration of other ions in cerebrospinal fluid 144 NaCl, 5.3 KCl, 2.5 CaCl_2_, 1 MgCl_2_, 1 NaH_2_PO_4_, 10 HEPES, 10 mM glucose, pH 7.4 (Agathou and Káradóttir, 2019). Temperature was controlled using a TC-324C automatic temperature controller (Warner Instruments). The pipette solution contained (in mM): 130 K-gluconate, 4 NaCl, 0.5 CaCl_2_, 10 HEPES 0.5 EGTA, pH 7.2 with KOH. Osmolarity of the external solution was 312 mOsm and the internal solution was 270 mOsm, both adjusted with sucrose. Borosilicate glass pipettes were pulled to a resistance of 5-20 MΩ.

To record from neurons, electrodes were brought into contact with the cell membrane and gigaOhm seals formed by gentle suction supplied from a syringe and water filled U tube. Firm suction was then used to break the membrane under the pipette to form a whole-cell recording. Recordings exhibiting a leak of more than 100 pA were rejected at this stage. Within 2 seconds of whole-cell access, the amplifier was switched to current clamp mode to enable instantaneous membrane potential to be estimated. Recordings were made for 2 minutes in current clamp mode to measure neuronal activity at baseline and after 5 minutes treatment with neuromodulators.

Action potential firing frequency was calculated using the frequency analysis function in WinEDR (Strathclyde electrophysiology software, http://spider.science.strath.ac.uk/sipbs/software_ses.htm). Data were presented as the number of action potentials per second (Hz). Action potentials were defined as regenerative, all-or-nothing changes in membrane potential that produced an overshoot beyond 0 mV.

### Statistical Analysis

Data were analysed using GraphPad Prism Version 10, with significance set at p ≤ 0.05. Statistical tests are indicated in the relevant Fig. legends.

## Supplemental Information

Figs 1, 2, 4-S6 and Tables 1 and 2

Supplementary Video 1, ‘Mbp mRNA in nascent sheaths in vitro’; related to Fig. 1H;

Supplementary Video 2, ‘Myelin peroxisome in internodal myelin’; related to Fig. 2

Supplementary Video 3, ‘Myelin peroxisome entering paranodal loop’; related to Fig. 2

Supplementary Video 4, ‘Myelin peroxisome traversing multiple paranodal loops’; related to Fig. 2

Supplementary Video 5, ‘Zoom of myelin peroxisome traversing paranodal loops’; related to Fig. 2

Supplementary Video 6, ‘β−tubulin IV in a tdTomato-labelled myelinating oligodendrocyte’; related to Fig. 2.

Supplementary Video 7, ‘Oligodendrocyte peroxisomes in the soma and processes’; related to Fig. 4

## Data and Code Availability section

Raw data will be published on a public database if the manuscript is accepted for publication.

