## Supplementary Figures for "A myelinic channel system for motor-driven organelle transport"

**Supplementary Table 1A.** Motor proteins identified by label-free mass spectrometry in myelin purified from the brains of c57Bl/6N mice (MM) according to Jahn et al. (2020) or in myelin purified from the white matter of humans (HS) according to Gargareta et al. (2022).

| **Motor protein** | **Akronym** | **MM** | **HS** | **Function** |
| --- | --- | --- | --- | --- |
| Dynein 1 heavy chain | DYNC1H1 | 1 | 1 | Motor protein |
| Dynein 1 intermediate chain 1 | DYNC1I1 | 0 | 1 | Motor protein |
| Dynein 1 intermediate chain 2 | DYNC1I2 | 1 | 1 | Motor protein |
| Dynein 1 light intermediate chain 1 | DYNC1LI1 | 1 | 1 | Motor protein |
| Dynein 1 light intermediate chain 2 | DYNC1LI2 | 1 | 1 | Motor protein |
| Dynein light chain 1 | DYNLL1 | 1 | 1 | Motor protein |
| Dynein light chain 2 | DYNLL2 | 1 | 1 | Motor protein |
| Kinesin heavy chain 5A | KIF5A | 0 | 1 | Motor protein |
| Kinesin heavy chain 5B | KIF5B | 1 | 1 | Motor protein |
| Kinesin heavy chain 5C | KIF5C | 1 | 1 | Motor protein |
| Kinesin light chain 1 | KLC1 | 1 | 1 | Motor protein |
| Kinesin light chain 2 | KLC2 | 0 | 1 | Motor protein |
| Kinesin-like protein 2A | KIF2A | 0 | 1 | Motor protein |
| Kinesin-like protein 21A | KIF21A | 0 | 1 | Motor protein |
| Myosin 1d | MYO1D | 1 | 1 | Motor protein |
| Myosin 1e | MYO1E | 1 | 0 | Motor protein |
| Myosin 5a | MYO5A | 1 | 1 | Motor protein |
| Myosin 5d | MYO5D | 0 | 1 | Motor protein |
| Myosin 6 | MYO6 | 1 | 0 | Motor protein |
| Myosin 18a | MYO18A | 1 | 1 | Motor protein |
| Myosin heavy chain 9 | MYH9 | 1 | 1 | Motor protein |
| Myosin heavy chain 10 | MYH10 | 1 | 1 | Motor protein |
| Myosin heavy chain 11 | MYH11 | 0 | 1 | Motor protein |
| Myosin heavy chain 14 | MYH14 | 1 | 1 | Motor protein |
| Myosin light chain 6 | MYL6 | 1 | 1 | Motor protein |
| Myosin light chain 6b | MYL6B | 0 | 1 | Motor protein |
| Myosin light chain 12b | MYL12B | 1 | 0 | Motor protein |

| **Supplementary Table 1B.** Selected proteins identified by label-free mass spectrometry in myelin purified from the brains of c57Bl/6N mice (MM) according to Jahn et al. (2020) or in myelin purified from the white matter of humans (HS) according to Gargareta et al. (2022). | | | | |
| --- | --- | --- | --- | --- |
| **Tubulin/actin associated proteins** | **Akronym** | **MM** | **HS** | **Function** |
| Centractin alpha | ACTR1A | 1 | 1 | Binds microtubules |
| Centractin beta | ACTR1B | 1 | 1 | Binds microtubules |
| Cofilin 1 | CFL1 | 1 | 1 | Actin modulating |
| Cofilin 2 | CFL2 | 1 | 1 | Actin modulating |
| Cyclic nucleotide phosphodiesterase | CNP | 1 | 1 | Binds microtubules |
| Dynactin subunit 1 | DCTN1 | 1 | 1 | Binds microtubules |
| Dynactin subunit 2 | DCTN2 | 1 | 1 | Binds microtubules |
| Dynactin subunit 3 | DCTN3 | 0 | 1 | Binds microtubules |
| Dynactin subunit 4 | DCTN4 | 1 | 1 | Binds microtubules |
| Histone deacetylase 11 | HDAC11 | 1 | 1 | Tubulin-modifying |
| Microtubule-associated protein 1A | MAP1A | 1 | 1 | Tubulin-modifying |
| Microtubule-associated protein 1B | MAP1B | 1 | 1 | Tubulin-modifying |
| Microtubule-associated protein 2 | MAP2 | 1 | 0 | Tubulin-modifying |
| Microtubule-associated protein 4 | MAP4 | 1 | 1 | Tubulin-modifying |
| Microtubule-associated protein 6 | MAP6 | 1 | 1 | Tubulin-modifying |
| Sirtuin 2 | SIRT2 | 1 | 1 | Tubulin-modifying |
| Tubulin alpha | TUBA | 1 | 1 | Tubulin |
| Tubulin beta | TUBB | 1 | 1 | Tubulin |
| Tubulin polymerization-promoting protein | TPPP | 1 | 1 | Tubulin-modifying |
| Tubulin polymerization-promoting protein 3 | TPPP3 | 1 | 1 | Tubulin-modifying |

**Supplementary Table 2. *Mbp* and *Mobp* probes used for smFISH**

| *Mbp* probe sequences | | Mobp probe sequences | |
| --- | --- | --- | --- |
| mMbp-exon_01 | gcctgtctttgaaggtgt | mMobp-exon_01 | aggttcctgtggatagcg |
| mMbp-exon_02 | gaagctcgtcggactctg | mMobp-exon_02 | gcgagtaatggctgttgt |
| mMbp-exon_03 | tggggtcttcttggatgg | mMobp-exon_03 | ggcaaccagaactctggt |
| mMbp-exon_04 | gtgagggtctcttctgtg | mMobp-exon_04 | ctgggttttcatccgtgc |
| mMbp-exon_05 | tggccaggtacttggatc | mMobp-exon_05 | ccttggccattttctgac |
| mMbp-exon_06 | gcatggtccatggtactt | mMobp-exon_06 | tggttcttggagagcctg |
| mMbp-exon_07 | ctgtgccttgggaggaag | mMobp-exon_07 | tgctgaagtgctcggaga |
| mMbp-exon_08 | aagaagcgcccgatggag | mMobp-exon_08 | tgaagggtgggcagcagt |
| mMbp-exon_09 | ctgctttagccagggtac | mMobp-exon_09 | cacgcttggagttgagga |
| mMbp-exon_10 | gcatgagagggcagaggg | mMobp-exon_10 | tgtacttgcggtccacga |
| mMbp-exon_11 | tgtacatgtggcacagcc | mMobp-exon_11 | aagcaaccgctcttgcag |
| mMbp-exon_12 | cagggagccataatgggt | mMobp-exon_12 | tagtcttctggcaggcac |
| mMbp-exon_13 | gttttcatcttgggtccg | mMobp-exon_13 | tctgaggggacgtggcac |
| mMbp-exon_14 | gaggtggtgttcgaggtg | mMobp-exon_14 | cagctggctggtgcttgg |
| mMbp-exon_15 | atctgctgagggacaggc | mMobp-exon_15 | ctctgaccaccactgggg |
| mMbp-exon_16 | tagccaaatcctggcttc | mMobp-exon_16 | ggactttggcttggctgg |
| mMbp-exon_17 | atagtcggaagctctgcc | mMobp-exon_17 | tggcttggctggcatcag |
| mMbp-exon_18 | ttgaatcccttgtgagcc | mMobp-exon_18 | attggcacagacacagc |
| mMbp-exon_19 | tttggaaagcgtgccctg | mMobp-exon_19 | ttagtgacttccagtcct |
| mMbp-exon_20 | atgggagatccagagcgg | mMobp-exon_20 | ggtggttaaggaggctca |
| mMbp-exon_21 | gagggcaggattcgggaa | mMobp-exon_21 | ctatgtctctctgaggca |
| mMbp-exon_22 | gggtctgctctaactagc | mMobp-exon_22 | aaatcgagcatcgggtcc |
| mMbp-exon_23 | cggctccacgggattaag | mMobp-exon_23 | agatgcgtcaggagggtc |
| mMbp-exon_24 | gttttaaccagtcggggt | mMobp-exon_24 | gtttgctgagtggctgta |
| mMbp-exon_25 | gtcagggctgagaagacc | mMobp-exon_25 | tccccagtacagcatatg |
| mMbp-exon_26 | acctccggagtcgaacaa | mMobp-exon_26 | gtctctttttggtactca |
| mMbp-exon_27 | cttggaagggtgtccgtg | mMobp-exon_27 | acagtacatcagggccag |
| mMbp-exon_28 | acggaagctggaggggtg | mMobp-exon_28 | gcatggcagtactggagt |
| mMbp-exon_29 | ccaagactgtctgatcct | mMobp-exon_29 | ccatgggcataaacacc |
| mMbp-exon_30 | aagcccccgttgtataag | mMobp-exon_30 | gctggaaagttcccagtt |
| mMbp-exon_31 | ccacgctcgaaatcagct | mMobp-exon_31 | ttctccagtttgctcact |
| mMbp-exon_32 | cagcgtgttctcctaagt | mMobp-exon_32 | taagttcctctctggcca |
| mMbp-exon_33 | atttggcggccacacata | mMobp-exon_33 | acccgaggtagagcagat |
| mMbp-exon_34 | ggagccactgaggagaga | mMobp-exon_34 | acttcttccttggggttg |
| mMbp-exon_35 | cctgggctctgagaggaa | mMobp-exon_35 | caagaagtcccttctcgg |
| mMbp-exon_36 | caccatgagaagtggcca | mMobp-exon_36 | caggcagaggctgtccat |
| mMbp-exon_37 | tgtaccaatgggcactgc | mMobp-exon_37 | ttgcgggaagcagtgagt |
| mMbp-exon_38 | ctttctaggaggcaggga | mMobp-exon_38 | attcgtctgagagcagat |
| mMbp-exon_39 | cccccctaaagctaagaa | mMobp-exon_39 | cctgcatcatacttgagg |
| mMbp-exon_40 | gcagcgactcgattcagt | mMobp-exon_40 | ttacatcagcaggcgtcc |
| mMbp-exon_41 | gtgtggggctctttggaa | mMobp-exon_41 | gactgagggcatctgtgg |
| mMbp-exon_42 | acatcaaccatcacctgc | mMobp-exon_42 | ctaaggctcccacaccat |
| mMbp-exon_43 | acccacactctactcaga | mMobp-exon_43 | tctcacctccaggaaggc |
| mMbp-exon_44 | accctcacgttattgtgg | mMobp-exon_44 | gtacgacacgaagagcc |
| mMbp-exon_45 | aaacactcccgtgggaca | mMobp-exon_45 | ggctccagatgaaacca |
| mMbp-exon_46 | ggtcgttcagtcacactg | mMobp-exon_46 | tgcacctgccagagagaa |
| mMbp-exon_47 | gtctggacgaagccatgg | mMobp-exon_47 | ttgccaaagaccctgagg |
| mMbp-exon_48 | tagtaggtgcttctgtcc | mMobp-exon_48 | gtgtggattagctctgca |

**Supplementary Figures**

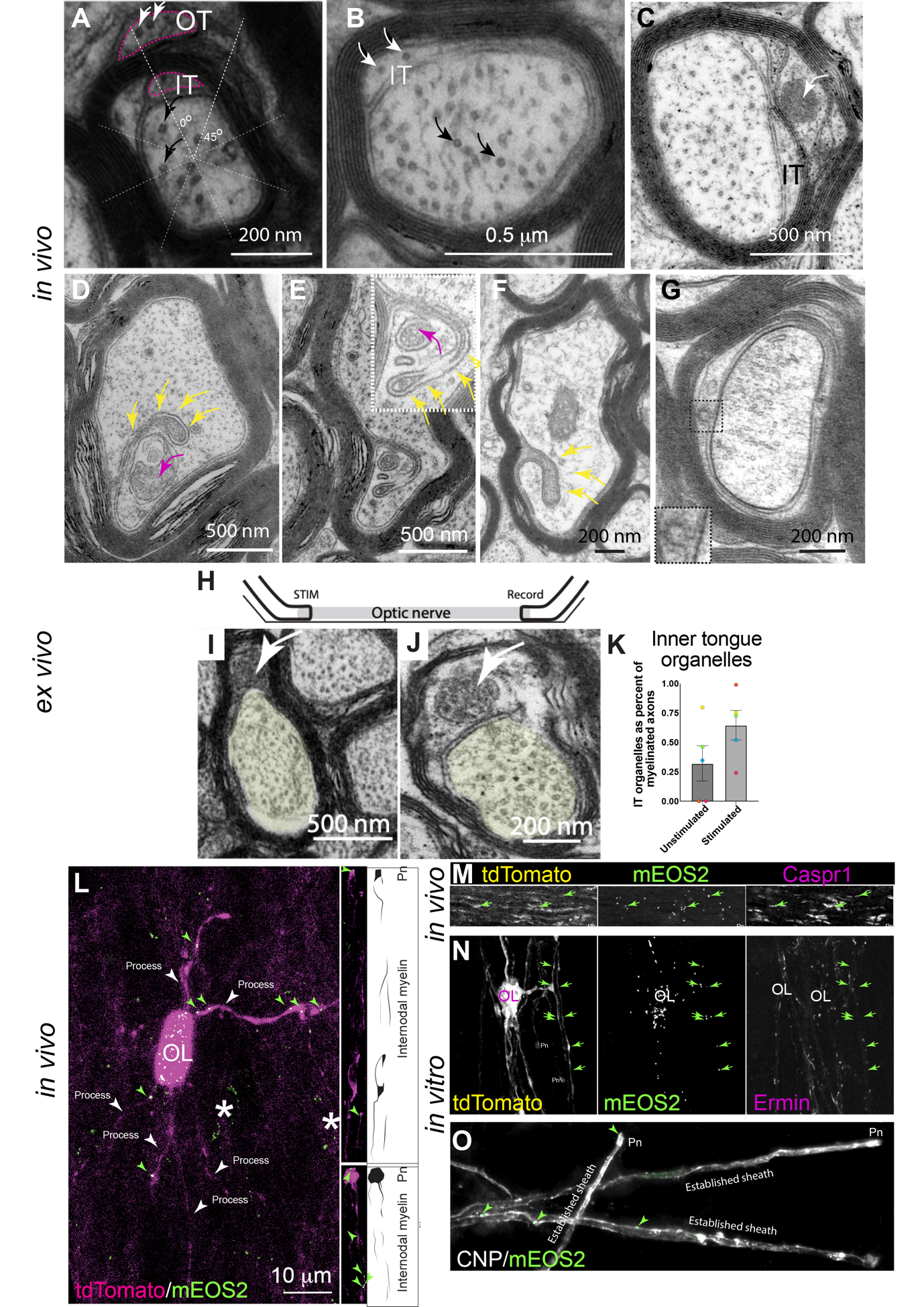

**Supplementary Figure 1. Microtubules and organelles populate the myelinic channel system.** Electron micrographs of optic nerve fibres of wild type mice. **A.** Microtubules (white arrows) in the outer tongue (OT) of myelin appear identical to those in the axon (black arrows). Here, the inner tongue (IT) and OT occur in the same octant, as delineated by dotted lines and reported previously (Peters, 1964). **B.** Microtubules (white arrows) in the IT (white arrows) and in the axon for comparison (black arrow). **C.** A dense organelle (white arrow) resembling a peroxisome is located between otherwise compacted layers of myelin. **D-F.** Glial invaginations of the axon (yellow arrows) often (D and E), but not always (F) appear to envelop axonal organelles (magenta arrows). **G.** A fusion profile on the glial membrane (black square and inset) that appears to open into the periaxonal space. **H.** One of each pair of adult mouse optic nerves were held between stimulating and recording suction electrodes and stimulated at 1 Hz for one minute then 7 (blue symbols in K) or 50 Hz (pink, orange, green and yellow symbols in K) for 20 minutes. The contralateral nerve was unstimulated. **I** and **J**. Myelin organelles in myelin’s inner tongue (white arrows). **K.** On average, there was an increase in dense organelles (peroxisome-like structures as in I or multivesicular bodies as in J) in the stimulated nerve compared to the unstimulated one, although the differences were not significant. Bars represent mean $\pm$SD. Data were analysed using an unpaired, two-tailed Student’s t-test. p = 0.1364. **L.** tdTomato labelled oligodendrocyte from an imprint of adult murine spinal cord in which mEOS2-labelled peroxisomes (green arrowheads) can be observed in the cell body (asterisk), processes (white arrows) and in myelin sheaths (the last shown as insets on the right), where they were often located in paranodal loops (Pn). For clarity, schematics of the sheaths are shown alongside. **M.** Individual channels from a zoomed in region of the composite image in Figure 1G, showing tdTomato, mEOS2-labelled oligodendroglial peroxisomes and axonal Caspr1 in a longitudinal section of adult mouse optic nerve. **N.** Individual channels from composite Figure 1J, showing a tdTomato labelled myelinating oligodendrocyte, mEOS2-labelled oligodendroglial peroxisomes and Ermin, *in vitro*. **O.** In another myelinating cell culture, three CNP +ve myelin sheaths terminate in paranodal loops. mEOS2-labelled peroxisomes are present in internodal and paranodal (Pn) regions of the sheaths.

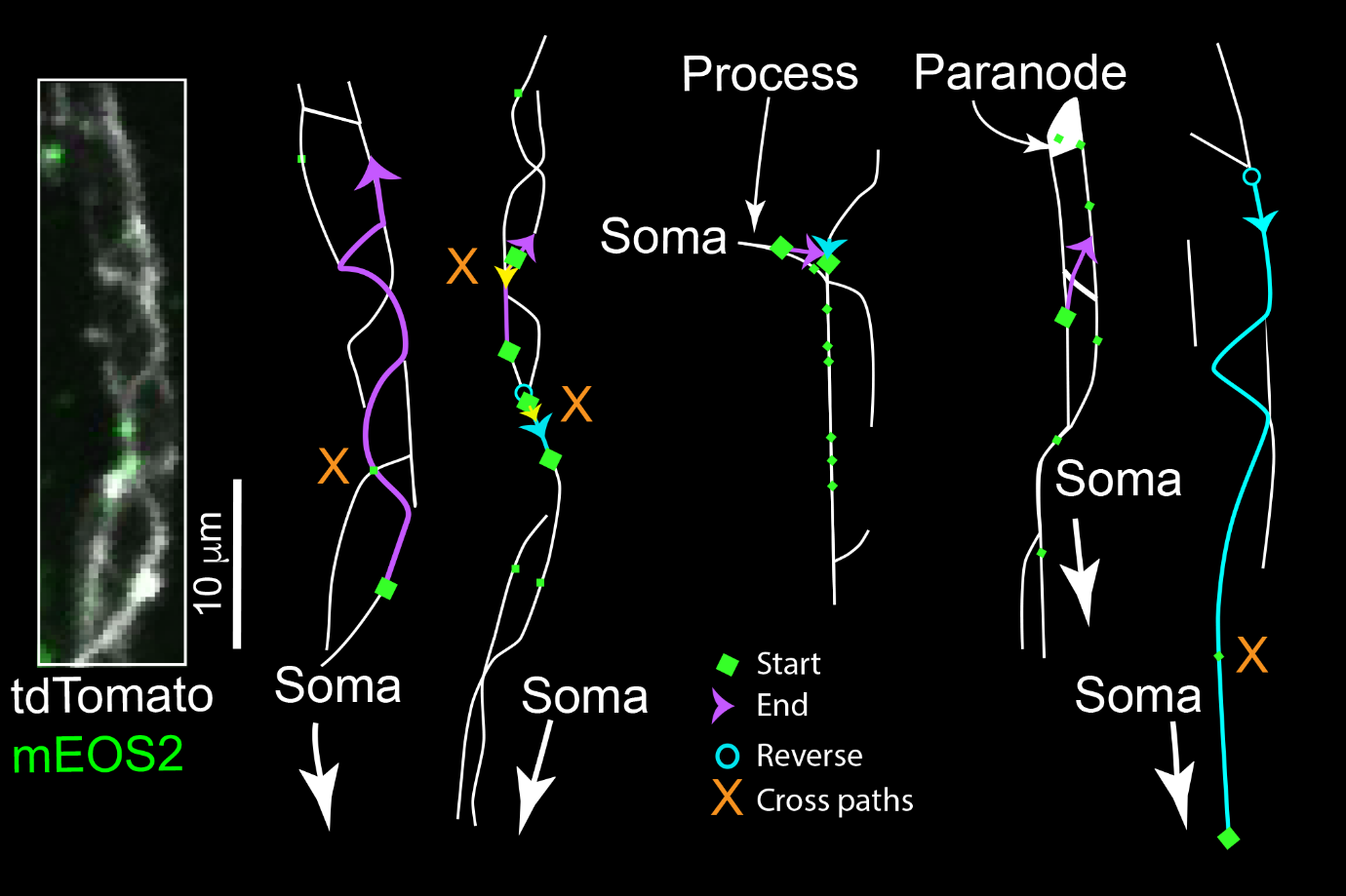

**Supplementary Figure 2A. Myelinic peroxisomes are motile.** Traces of tdTomato rendered sheaths (white) and peroxisomes within them (green diamonds) from 10-minute time-lapse images from a single z-plane using a spinning disc confocal microscope. On the left, the inset shows a maximum intensity projection (MIP) of a confocal z-stack of the sheath illustrated on its right. Peroxisomes whose paths are shown are indicated by a large green diamond at the start of the movement; those that remain stationary, are indicated by small green diamonds. Magenta arrows show the paths of peroxisomes away from the cell body; yellow arrows show paths to the cell body; and cyan arrows show paths taken by peroxisomes that reverse direction. Arrowheads represent the final destinations within the timeframe. Motile peroxisomes often cross paths with stationary ones (orange Xs) or reverse their direction of travel (cyan circles). See also Videos S2, S3, S4 and S5.

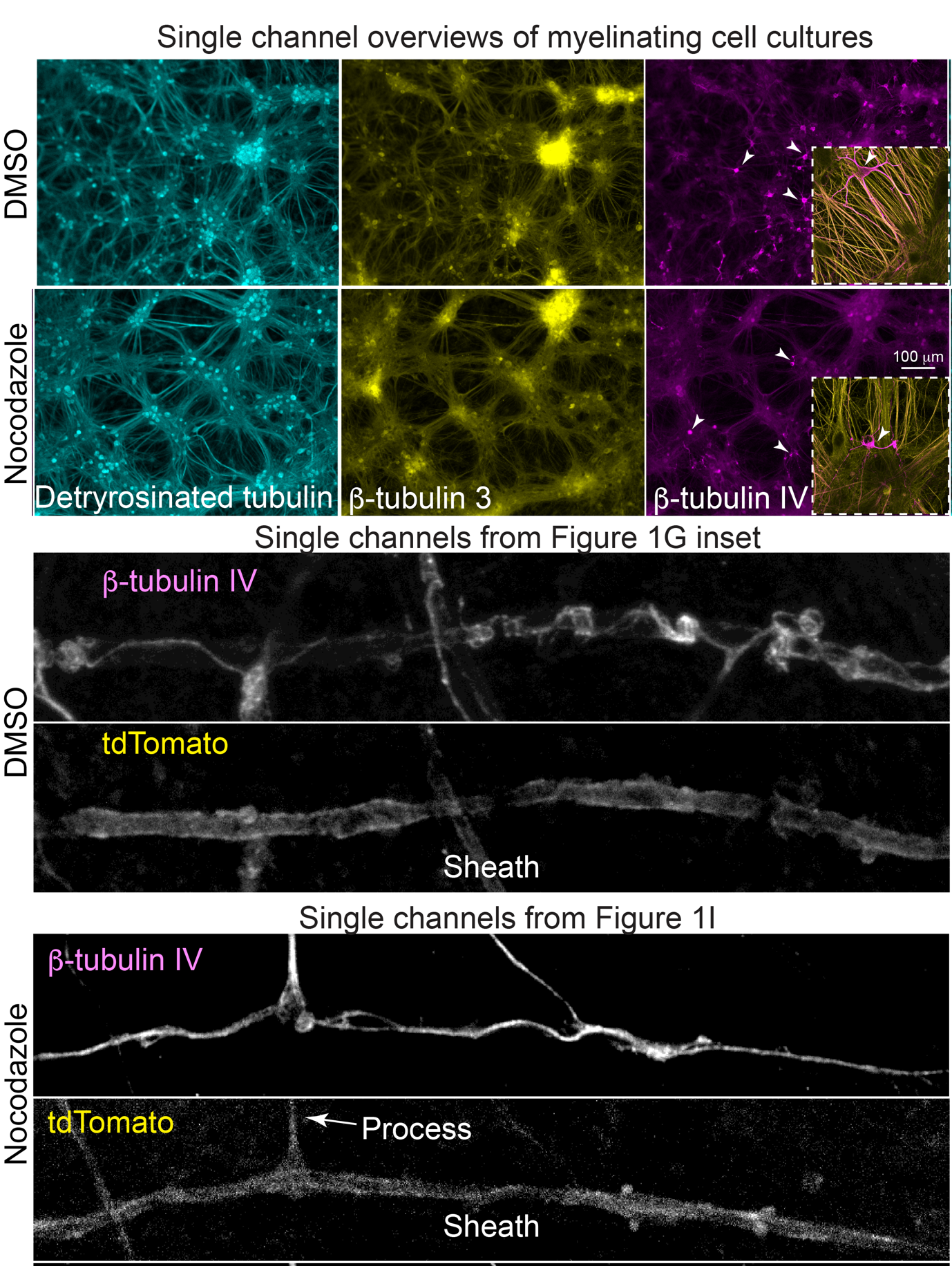

**Supplementary Figure 2B. Nocodazole partially impairs the microtubule polymer in myelinating cells cultures.** Fluorescence micrographs of microtubules in myelinating cell cultures treated with DMSO or nocodazole, following extraction of free tubulins. Cells were labelled with anti-detyrosinated tubulin (cyan), a marker of ‘stable’ microtubules; β-tubulin 3 (yellow), which is specific in the CNS to neurons and their processes; and β-tubulin 4 (magenta), which is specific in the CNS to oligodendrocytes (white arrowheads), including myelin sheaths. In these cell dense cultures, incubation with nocodazole does not fully perturb the microtubule polymer as can be observed in the images on the left of detyrosinated microtubules and the insets on the right where many β-tubulin 3 and 4 containing microtubules appear intact. Overall, however, β-tubulin 3 labelling appears less well-defined in the nocodazole treated cultures compared to the DMSO treated controls (middle panels and insets on the right) due to nocodazole’s interference with microtubule polymerisation.

The lower panels illustrate the individual channels from Figures 2G inset and 2I, demonstrating that many β−tubulin IV containing microtubules are resistant to nocodazole under the conditions used (20 µM, 3 hours) to treat these cell dense cultures.

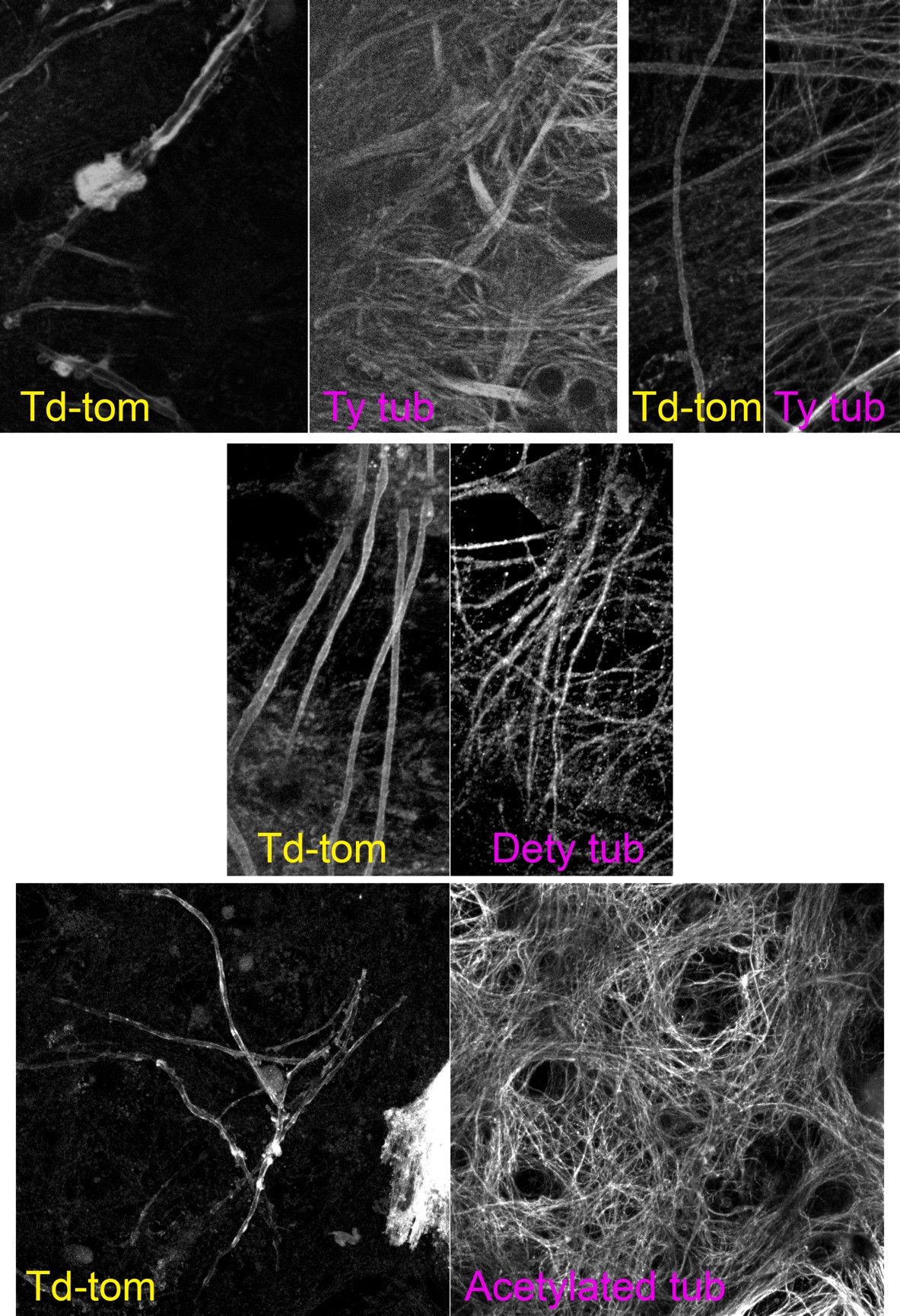

**Supplementary Figure 3. The myelinic channel system contains microtubules with various post-translational modifications.** Individual channels of the composite images shown in Figures 3C, D, E and F, showing tdTomato labelled myelinating oligodendrocytes and/or myelin sheaths, and microtubules in these and neighbouring cells. The tubulin extraction protocol, involving glutaraldehyde, causes background autofluorescence that can be seen particularly in the red channel (tdTomato).

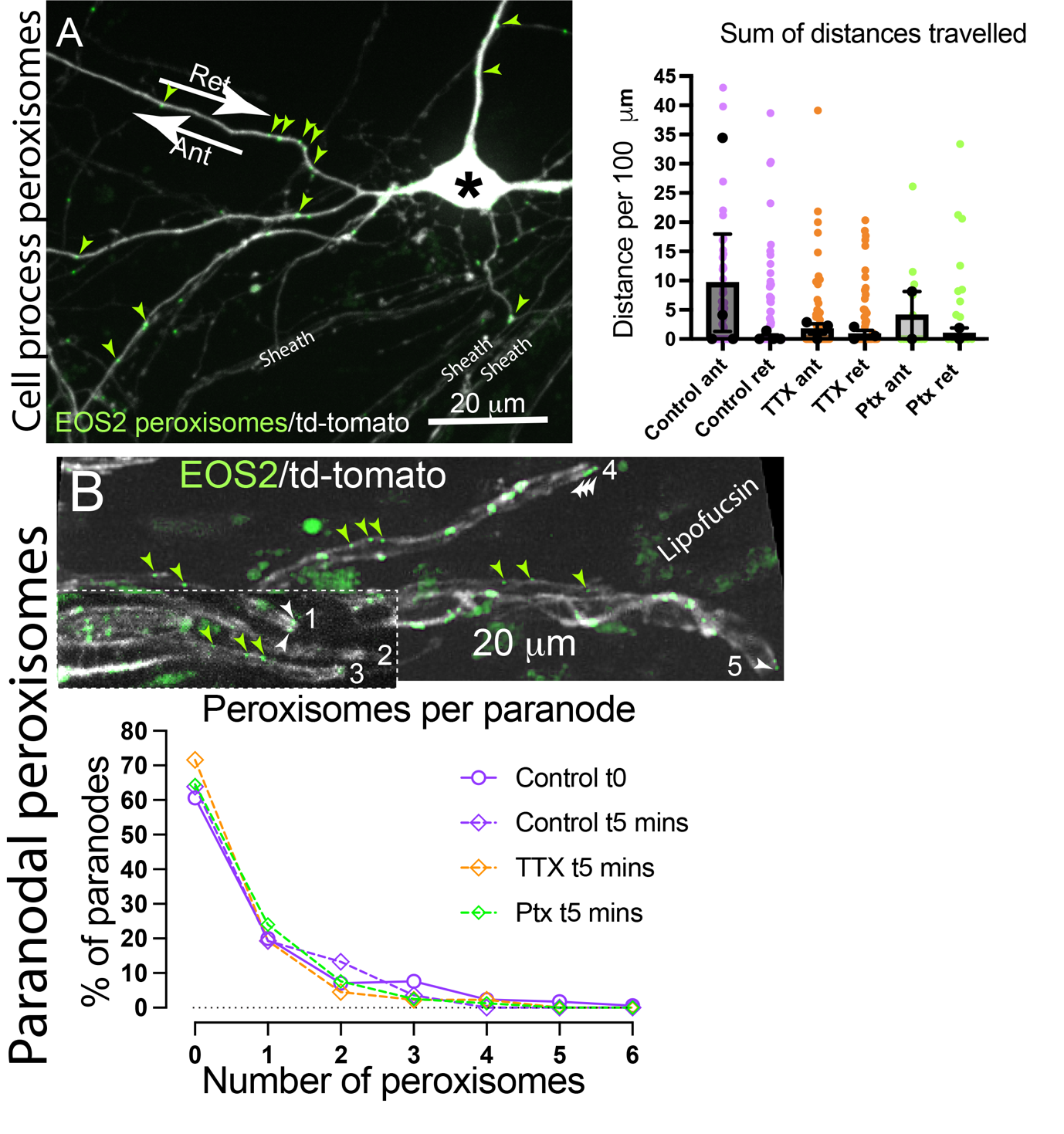
 **Supplementary Figure 4. Modulating neuronal activity does not modify peroxisome movement in processes or accumulation in the paranodes. A.** Movement of peroxisomes in processes, to (retrogradely) or from (anterogradely), the cell body is not altered by changes in neuronal activity. Coloured dots represent individual peroxisomes; black dots represent experimental medians. Bars represent the mean of the experimental medians ± SD. See also Video S6. **B.** Neuronal activity does not alter, within 5 minutes of drug application, the numbers of peroxisomes at paranodal regions of myelin. The images show two MIPs of sheaths with paranodal endings containing no (2), one (3), two (1 and 5), or three (4) peroxisomes. The graph shows the average (*n* = 4 independent experiments) percentage of paranodes with 0 to 6 peroxisomes per sheath.

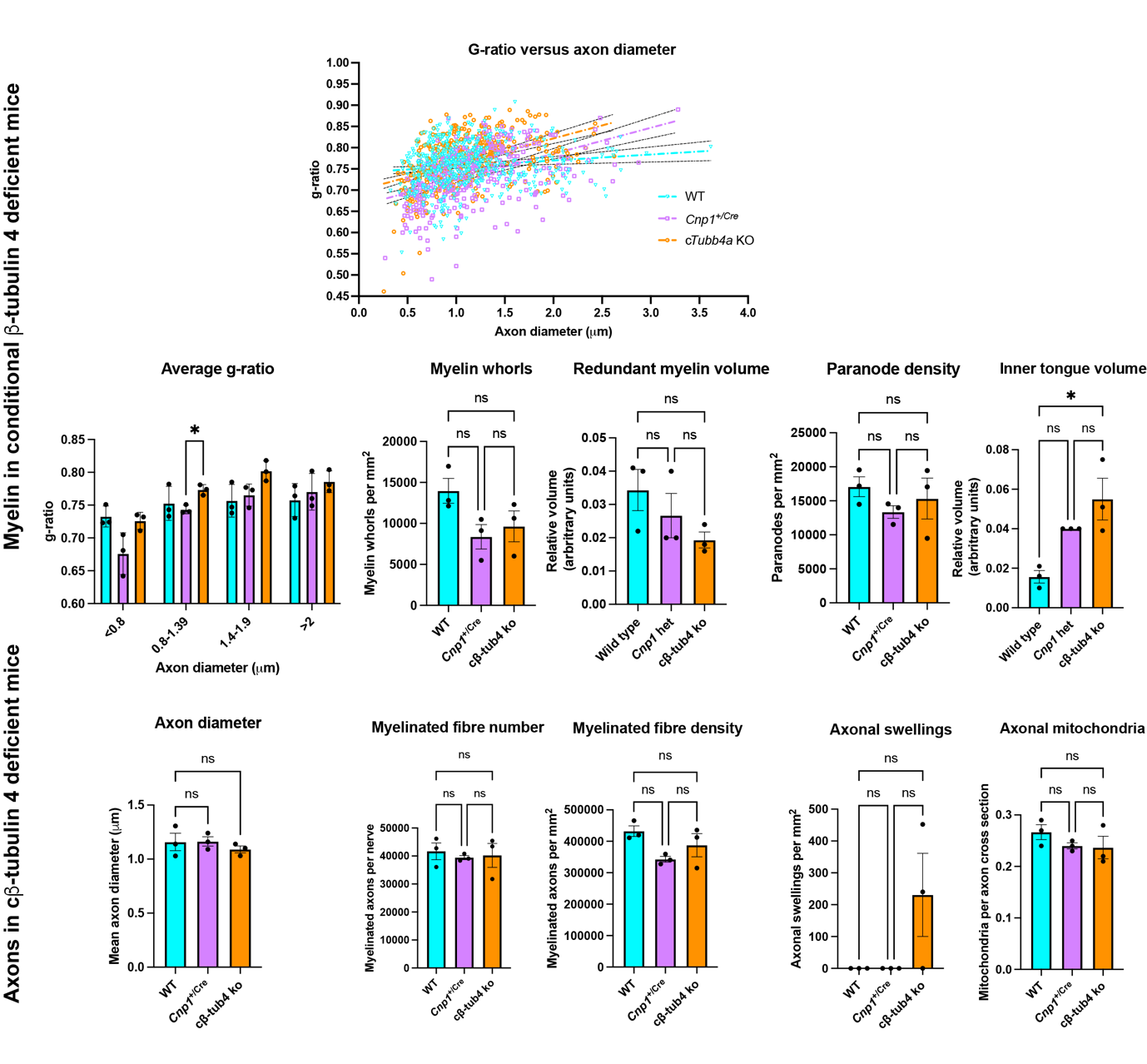

**Supplementary Figure 5A. Myelin and axons are largely unaltered in conditional β-tubulin 4 deficient mouse optic nerve.** Quantification of electron micrographs of optic nerve fibres of 14-month-old Tubb4a^flox/flox^;Cnp^+/+^ (WT), Tubb4a^+/+^;Cnp^+/Cre^ (Cnp1^+/Cre^) and Tubb4a^flox/flox^;Cnp^+/Cre^ (cβ-tub4 ko) showed that in axons larger than 0.8 mm diameter, myelin tended to be thinner in the conditional knockout (higher g-ratio) compared to controls, but the differences were mainly not significant. For g-ratio versus axon diameter, data were analysed using a 2-way ANOVA (p = 0.0303 for g-ratio x genotype) followed by a Tukey’s multiple comparisons test. For myelin whorls, redundant myelin, paranodes, inner tongue volume, axon number, axon density, axonal swellings, axonal mitochondria, data were analysed using an ordinary one-way ANOVA (p = 0.1128, p = 0.2280, p = 0.4688, p = 0.8698, p = 0.0993, p=0.0132, p = 0.0993, p = 0.1177. p = 0.3852, respectively) followed by a Tukey’s multiple comparisons test. The inner tongue volume was significantly higher in the cβ-tubulin IV ko than in the WT control but not significantly different from the *Cnp1*^Cre/+^ control suggesting CNP haploinsufficiency is the main contributor to inner tongue enlargement. * p < 0.05.

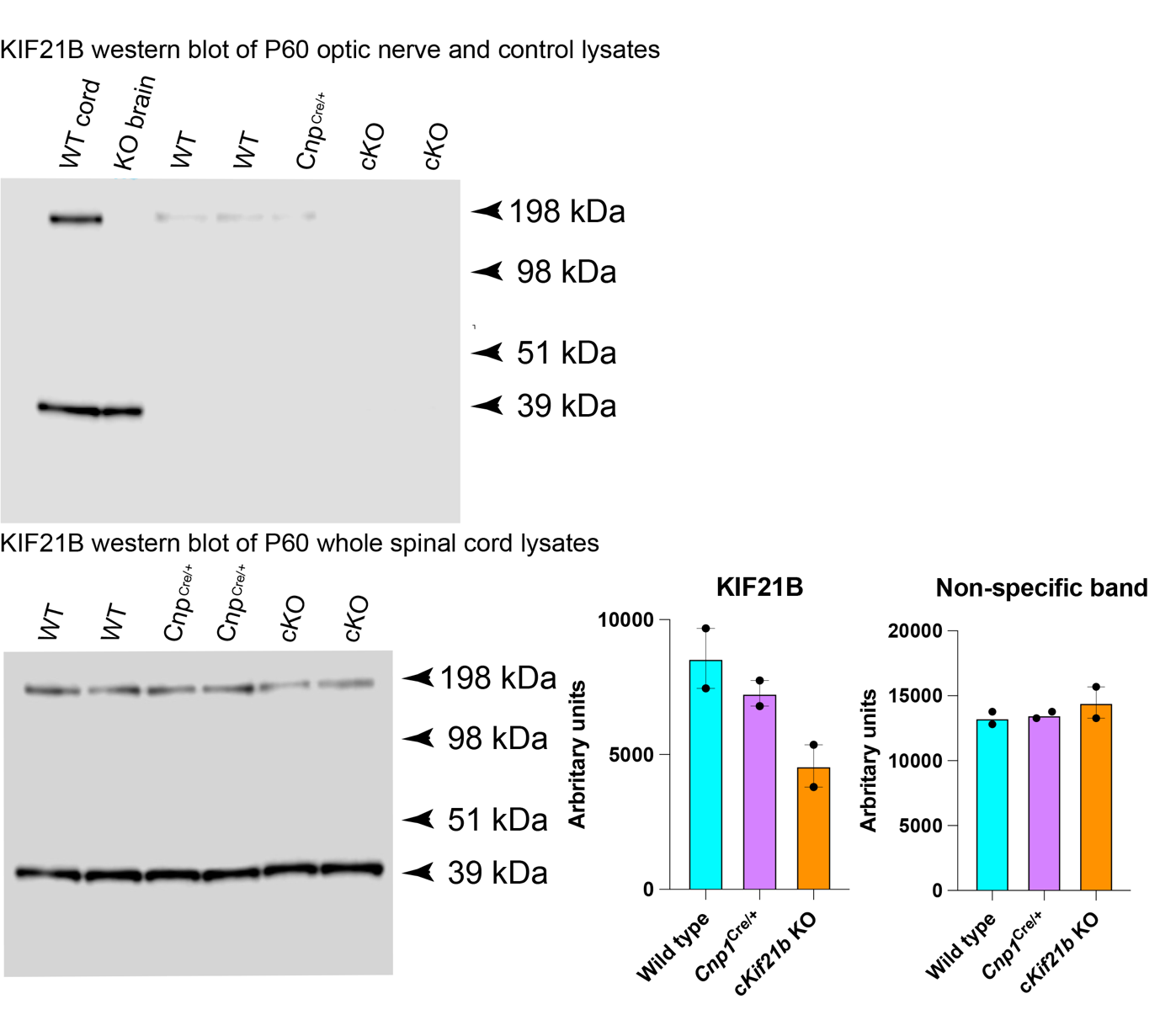
 **Supplementary Figure 5B. Western blotting of whole optic nerve and spinal cord lysates indicates Kif21b is enriched in oligodendrocytes, as well as neurons.** In western blots of optic nerve (right hand side of upper panel) which does not contain neuronal cell bodies or dendrites, Kif21b is markedly diminished in the *Kif21b* conditional knockout mice (cKO) in comparison to Kif21b^flox/flox^;Cnp^+/+^ (WT) and Kif21b^+/+^;Cnp^+/Cre^ (Cnp1^+/Cre^) controls. This indicates Kif21b is enriched in *Cnp1*-expressing cells (oligodendrocytes) in this white matter tract. Whole spinal cord lysate from a wild type mouse and brain from a global knockout mouse were run alongside to confirm antibody specificity. In western blots of whole spinal cord lysates (lower panel), Kif21b was only slightly diminished in the c*Kif21b* KO, as expected for a protein that is expressed in both neurons and oligodendrocytes. A band at ~39 kDa, due to secondary antibody binding non-specifically in cord and brain lysates, confirms equal loading of samples. Bars represent mean ± SD. Data points from individual animals.
