## Supplementary Videos for "A myelinic channel system for motor-driven organelle transport"

### Slide 1
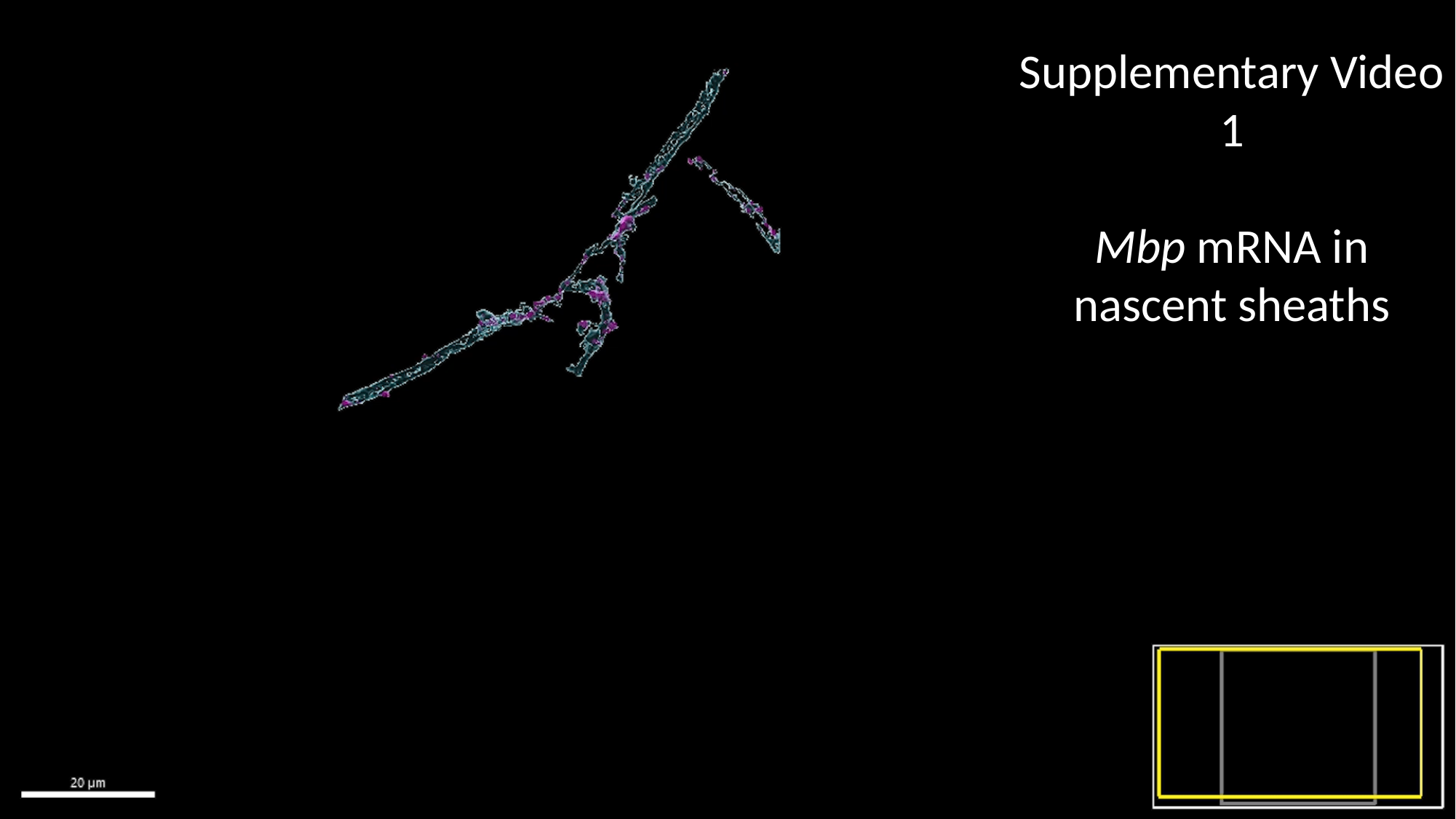

Supplementary Video 1
Mbp mRNA in nascent sheaths

### Slide 2
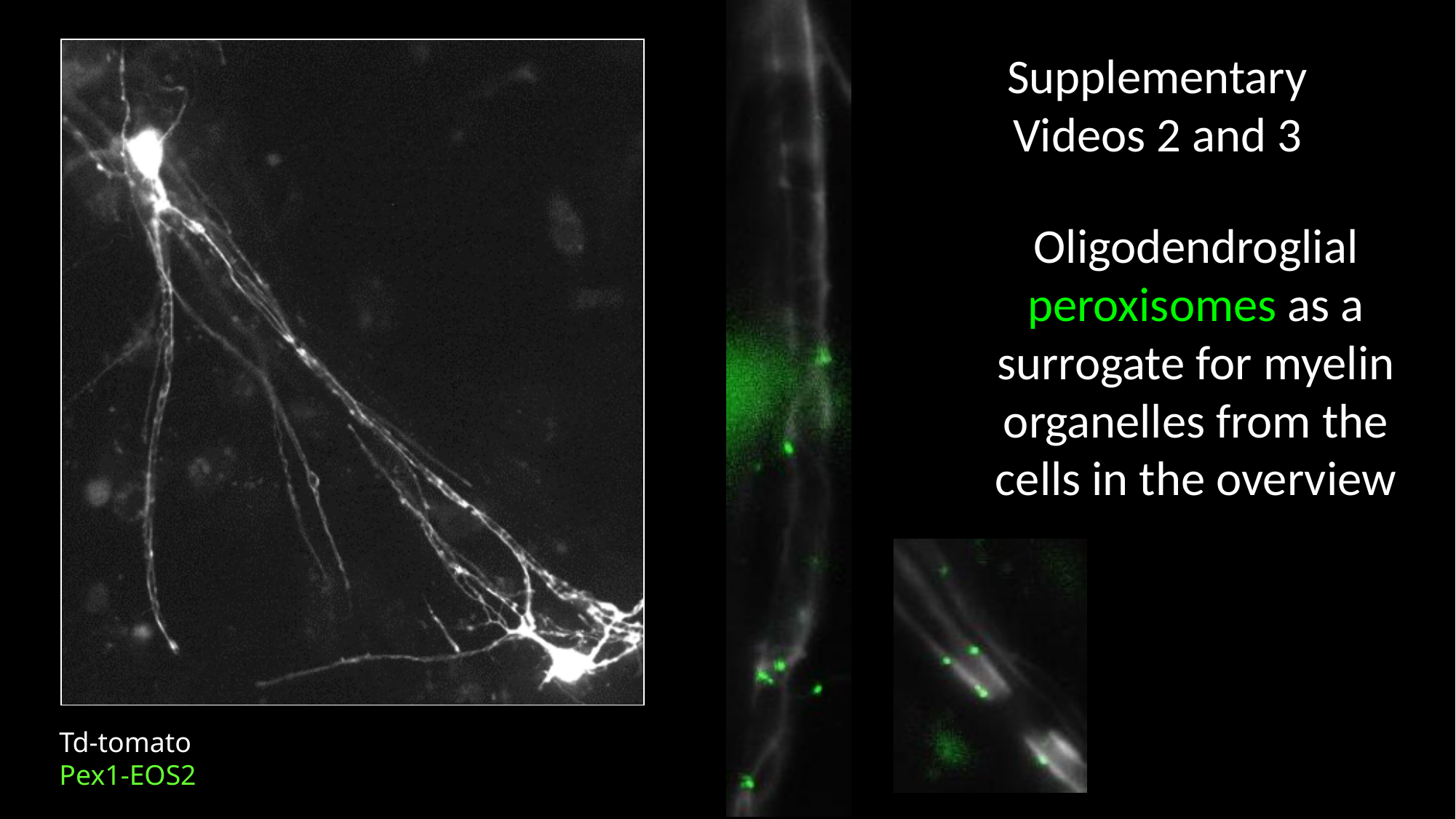

Supplementary Videos 2 and 3
Oligodendroglial peroxisomes as a surrogate for myelin organelles from the cells in the overview
Td-tomato
Pex1-EOS2

### Slide 3
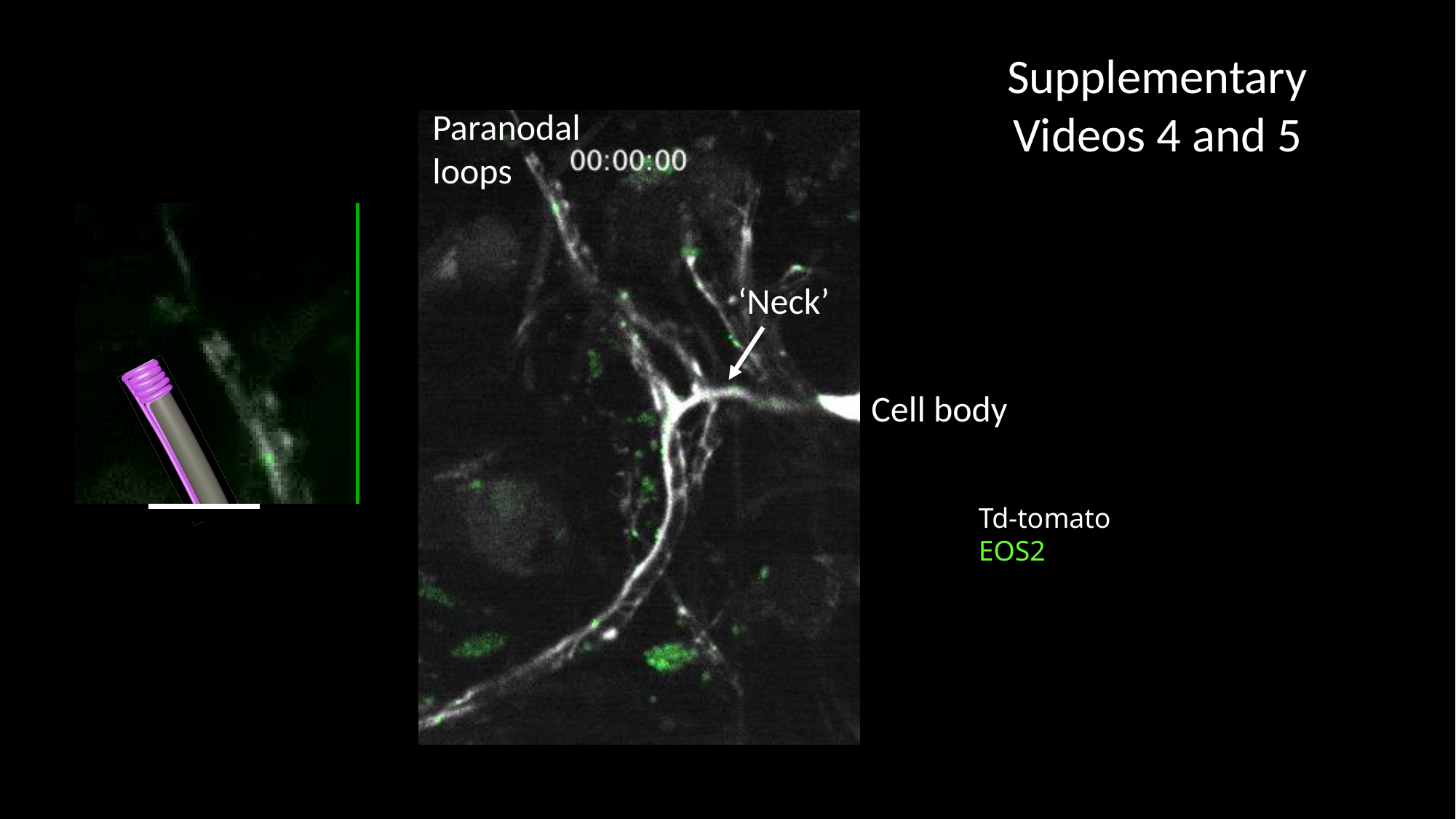

Supplementary Videos 4 and 5
Paranodal
loops
‘Neck’
Cell body
Td-tomato
EOS2

### Slide 4
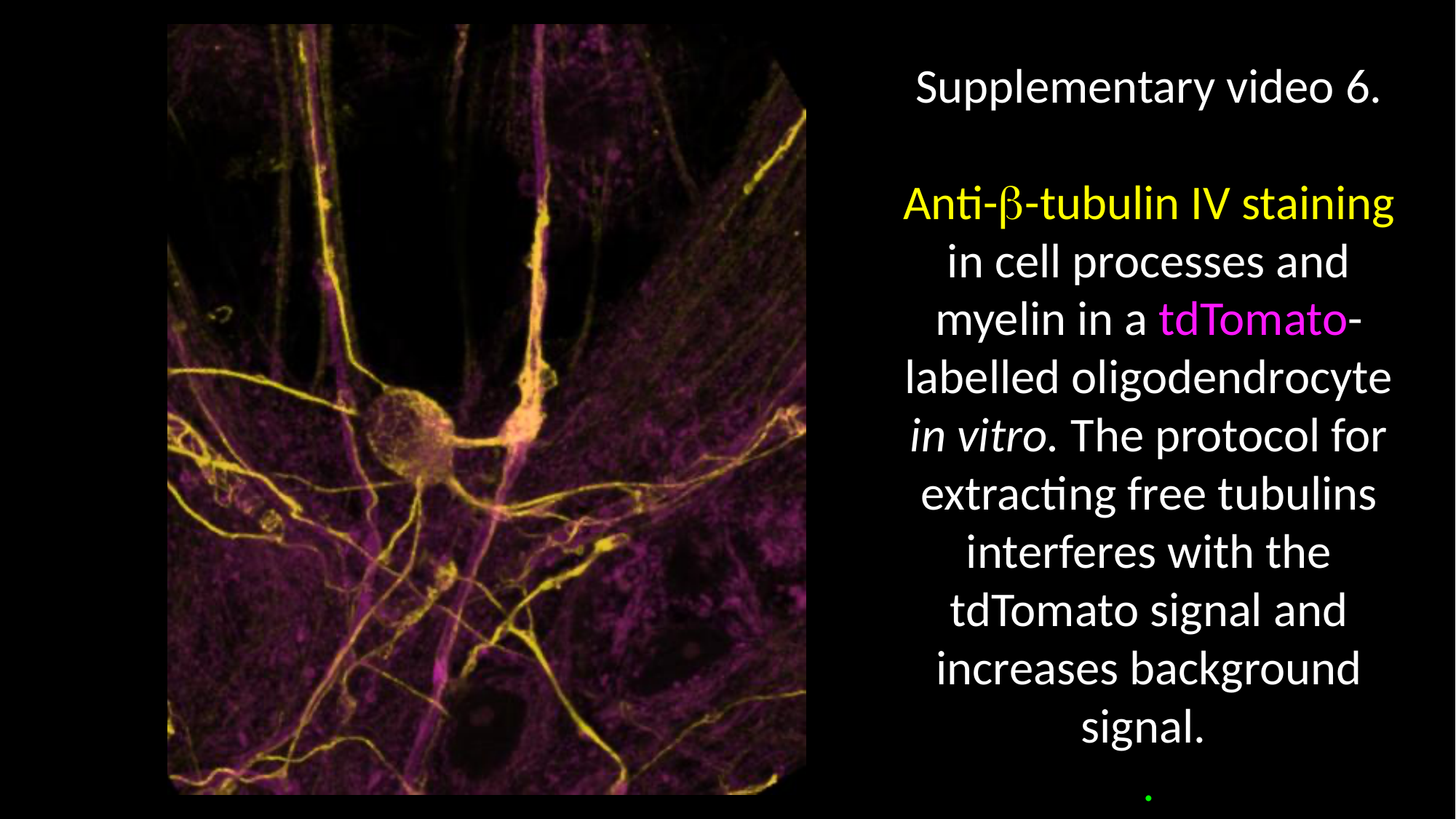

Supplementary video 6.
Anti-β-tubulin IV staining in cell processes and myelin in a tdTomato-labelled oligodendrocyte in vitro. The protocol for extracting free tubulins interferes with the tdTomato signal and increases background signal.
.

### Slide 5
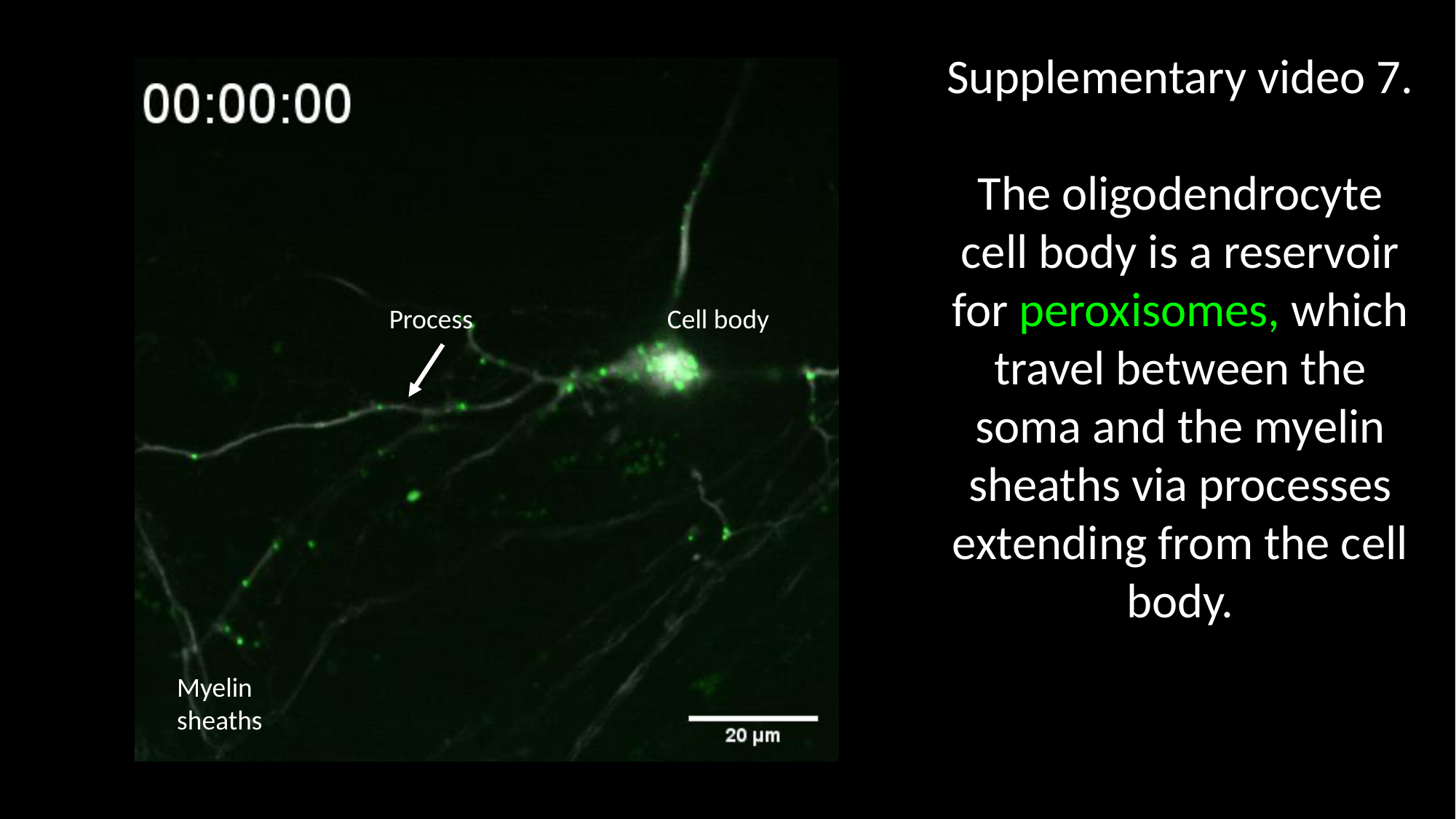

Supplementary video 7.
The oligodendrocyte cell body is a reservoir for peroxisomes, which travel between the soma and the myelin sheaths via processes extending from the cell body.
Process
Cell body
Myelin sheaths
